# Single-cell analysis of hepatoblastoma identifies distinct tumor cell signatures that predict susceptibility to chemotherapy using patient-specific tumor spheroids

**DOI:** 10.1101/2021.10.13.464268

**Authors:** Hanbing Song, Simon Bucher, Katherine Rosenberg, Margaret Tsui, Deviana Burhan, Soo-Jin Cho, Arun Rangaswami, Franklin W. Huang, Amar Nijagal, Bruce Wang

**Author notes:** These authors contributed equally as first authors (Hanbing Song, Simon Bucher). These authors contributed equally as senior authors who jointly supervised the work (Franklin W. Huang, Amar Nijagal, Bruce Wang).

## Abstract

Pediatric hepatoblastoma (HB) is the most common primary liver cancer in infants and children. Studies of HB that focus exclusively on tumor cells demonstrate sparse somatic mutations and a common cell of origin, the hepatoblast, across patients. In contrast to the homogeneity these studies would suggest, HB tumors have a high degree of heterogeneity that can portend poor prognosis. In this study, we used single-cell genomic techniques to analyze resected human pediatric HB specimens. This study establishes that tumor heterogeneity can be defined by the relative proportions of five distinct subtypes of tumor cells. Notably, patient-derived HB spheroid cultures predict differential responses to treatment based on the transcriptomic signature of each tumor, suggesting a path forward for precision oncology for these tumors. Collectively, these results define HB tumor heterogeneity with single-cell resolution and demonstrate that patient-derived spheroids can be used to evaluate responses to chemotherapy.

---

Hepatoblastoma (HB) is the most common primary pediatric liver cancer, accounting for approximately 1% of all pediatric malignancies, and its incidence is rising[1]. Five-year survival for HB is among the lowest for childhood cancers, driven by the 20% of cases that are chemotherapy resistant or unresectable[2, 3]. Current clinical risk stratification remains dependent on imaging and histological features at the time of diagnosis, with serum AFP as the only molecular marker[4]. There is an urgent need to improve the molecular characterization of HB to more accurately risk stratify patients. For patients with advanced HB, no effective treatment options exist outside of liver transplantation[5]. Progress in treating aggressive HB has been limited by the lack of models that reflect the heterogeneity of this tumor and that can be used to identify novel therapies[6].

The significant cellular heterogeneity observed in HB, both within and across patients[7], likely accounts for the limited utility of genomic studies from bulk tumor tissue for cancer staging[8–11]. Methods now exist for analyzing gene expression at the level of individual cells from dispersed neoplastic and normal tissues[12]. In this study we use single cell RNA sequencing (scRNA-seq) to distinguish HB tumor cells from non-tumor cells and to identify distinct tumor cell types that account for the heterogeneity of HB. We also use a novel method to grow HB tumor cells as patient-specific spheroids (PDS) and show how this can be used to predict treatment response and identify novel therapeutic targets.

## RESULTS

### Single-cell profiling of human pediatric HB reveals tumor-associated and patient-specific tumor cell populations

We established a workflow to isolate fresh pediatric HB tumor tissue at the time of surgical resection from nine patients (**Figure 1a**). Our samples included both epithelial and mixed epithelial mesenchymal tumors, each of which exhibited a range of epithelial histology, though none had small cell undifferentiated features indicating high risk (**Supplemental Table 1**). Eight of the nine patients underwent chemotherapy prior to resection. Single cells from tumor and paired adjacent normal tissues were isolated for scRNA-seq analysis. One of the nine patients had tumor extension into the adjacent tissue and therefore non-tumor tissue was unavailable. A total of 44,550 cells were captured, and 29,968 cells passed quality control (13,870 tumor, 16,098 non-tumor tissue) and were analyzed (**Supplemental Figure 1, Supplemental Table 2**). Using unbiased clustering and UMAP visualization, we identified 36 distinct clusters of cells (**Supplemental Figure 2**). We observed minimal inter-sample batch effect (**Supplemental Figure 3**).

**Figure 1.**
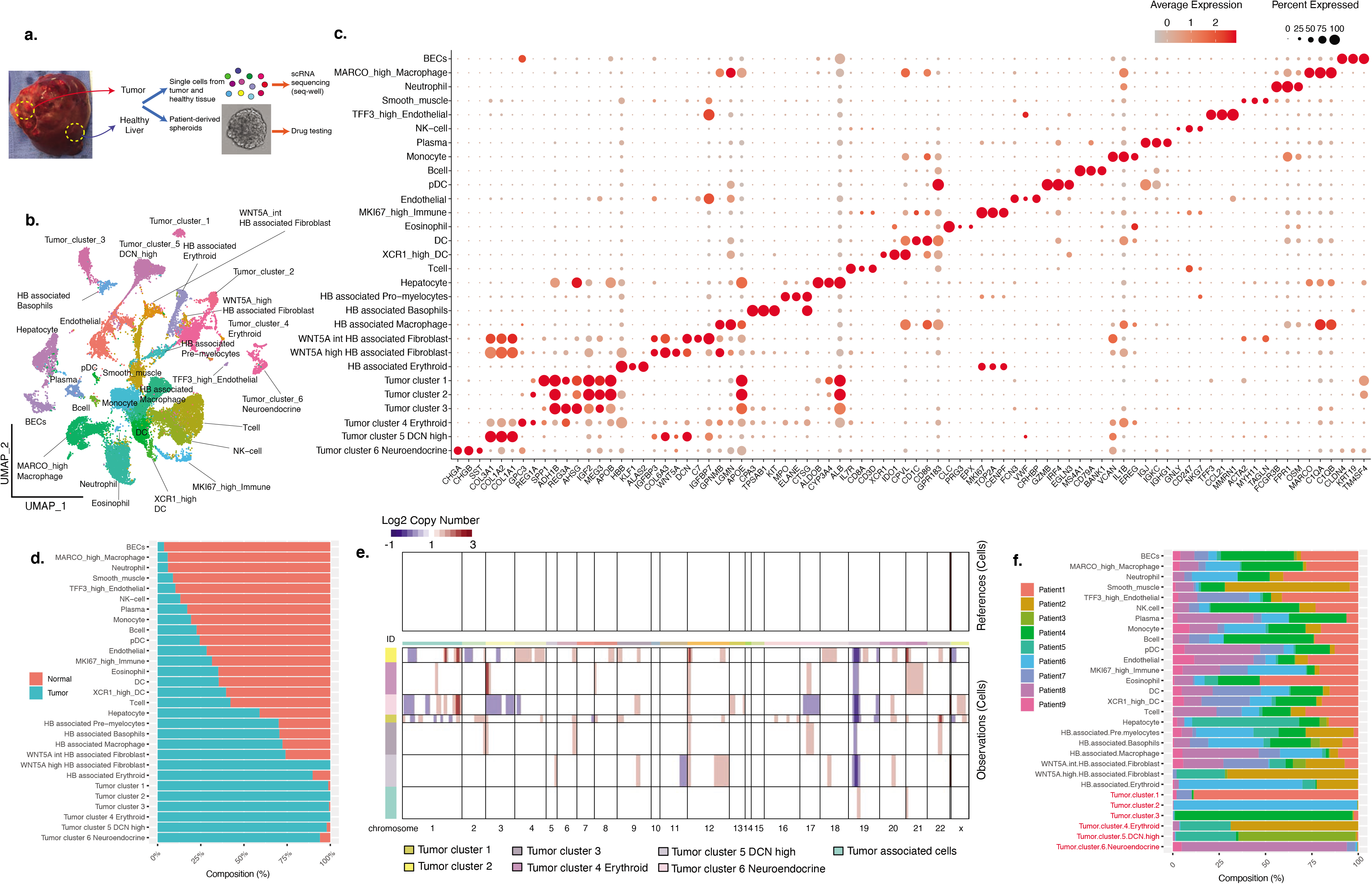
Single-cell profiling of HB patients revealed distinctive tumor cell populations and tumor associated populations. a. Flowchart of tissue processing of HB tumor and adjacent paired normal tissue samples for single-cell RNA sequencing and patient-derived spheroid culture. b. Uniform manifold approximation and projection (UMAP) of 29,968 cells from nine HB patients, annotated by cell types. c. Dot plot of all identified populations, each characterized by three known cell type markers. Average expression was indicated by the color gradient and the percentage of marker expressed was represented by the dot size. d. Stacked bar charts showing the contribution of the two sample types to each cell population (normal and tumor), ranked by the contribution from normal samples. e. Estimated copy-number alteration profile of all tumor and tumor-associated cell clusters using all the non-tumor and non-HB-associated cells as reference. Chromosomes are labeled on the horizontal axis. Estimated copy numbers are shown in blue (deletion) and red (amplification) color bars. f. Stacked bar charts show the contribution of the nine patients to each cell population.

To annotate these clusters, we performed differential gene expression analysis and identified 2,130 differentially expressed genes (DEGs) (log2 fold change > 1.0) (**Supplemental Table 2, Supplemental Figure 2**). We manually selected three well-recognized and highly significant DEGs for each cluster and applied a descriptive nomenclature, defining 29 cell types (**Figure 1b,c**). Seventeen of these were epithelial, stromal or immune cell populations that are normally resident in the liver. Epithelial populations include hepatocytes and biliary epithelial cells. Stromal cell populations include two clusters of endothelial cells (*VWF+/MMRN1+*) differentiated by the level of *TFF3* expression, and one cluster of *MYH11+/ACTA2+* smooth muscle cells. Immune cells identified include T-cells, NK-cells, monocytes, neutrophils, B-cells, plasma cells, eosinophils, macrophages, three clusters of dendritic cells (DC), and *MKI67* expressing proliferating immune cells (**Figure 1c**). We used an automated cell type annotation tool SingleR to validate our manual annotations (**Supplemental Figure 4)**[13].

In the remaining 12 clusters, >60% of cells originated from tumor suggesting these were either HB tumor cells or HB-associated cells (**Figure 1d, Supplemental Figure 3**). We designated six as “tumor cell clusters” based on their expression of high levels of known HB genes (**Figure 1c, Supplemental Figure 5, Supplemental Table 2**). Tumor cell clusters 1, 2, and 3 were characterized by high expression of the known HB tumor markers *REG3A, MEG3 and IGF2*[8]. Tumor cell cluster 4 expressed high levels of erythroid genes including *HBB, HBG1/2* and *ALAS2*. Tumor cell cluster 5 expressed high levels of fibroblast genes including *DCN*. Tumor cell cluster 6 expressed neuroendocrine markers (e.g. *CHGA* and *CHGB*).

The remaining six clusters did not express HB markers and were determined to be “HB-associated clusters”. They included three immune cell populations: a pre-myelocyte population, a tumor-associated macrophage (TAM) population showing low *MARCO* expression, and a basophil cluster. One HB-associated cluster (*HBB+/ALAS2+*) expressed high levels of erythroid progenitor genes (*KLF1*) and was highly proliferative, which we annotated as HB-associated erythroid cells [14]. We also identified two fibroblast populations differentiated by *WNT5A* expression level, which we annotated as *WNT5A*-intermediate and *WNT5A*-high HB-associated fibroblasts. While these two HB-associated clusters had gene expression profiles similar to that of the *DCN*-high tumor cell cluster 1, they did not express HB genes including *GPC3, DLK1* and *DKK1* (**Supplemental Figure 5**).

To validate the tumor cell annotations, we performed inferCNV analysis to compare the estimated copy number profiles for all tumor and tumor-associated populations in our dataset using the 17 nontumor associated populations as reference[15]. Compared to reference cells, only the six tumor cell clusters exhibited significantly different CNV profiles, supporting their identity as tumor cells (**Figure 1e**).

Interestingly, we found that four of the tumor cell clusters were unique to one patient while two were identified in two patients, indicating significant patient-specificity (**Figure 1f**). By contrast, all 17 nontumor-associated populations were present in every patient, with similar transcriptomes in all (**Figure 1f, Supplemental Figure 3**).

### Five distinct types of HB tumor cells account for the heterogeneity observed in HB tumor samples

We performed sub-cluster analysis to further characterize HB tumor cells, which yielded eight tumor cell populations (**Figure 2a, Supplemental Figure 6, Supplemental Table 3**). When investigating the composition of each population, we found that Patients 2 and 5 both contributed to the erythroid tumor cluster 6, consistent with the finding of extramedullary erythropoiesis in these two tumors (**Figure 2b, Supplemental Figure 6, Supplemental Figure 7**). Patients 3 and 5 both contributed to the *DCN*-high tumor cluster 7, consistent with fibrotic regions present in both tumors (**Supplemental Figure 7**). Closer examination of the DEGs showed that clusters 1-5 shared expression of known HB markers *GPC3, PEG10, REG3A, RELN*, and *PDK4* **(Supplemental Figure 6)**[8, 9, 16]. Notably, these populations are composed almost exclusively of cells from the four epithelial tumors (**Supplemental Table 1, Supplemental Figure 7**).

**Figure 2.**
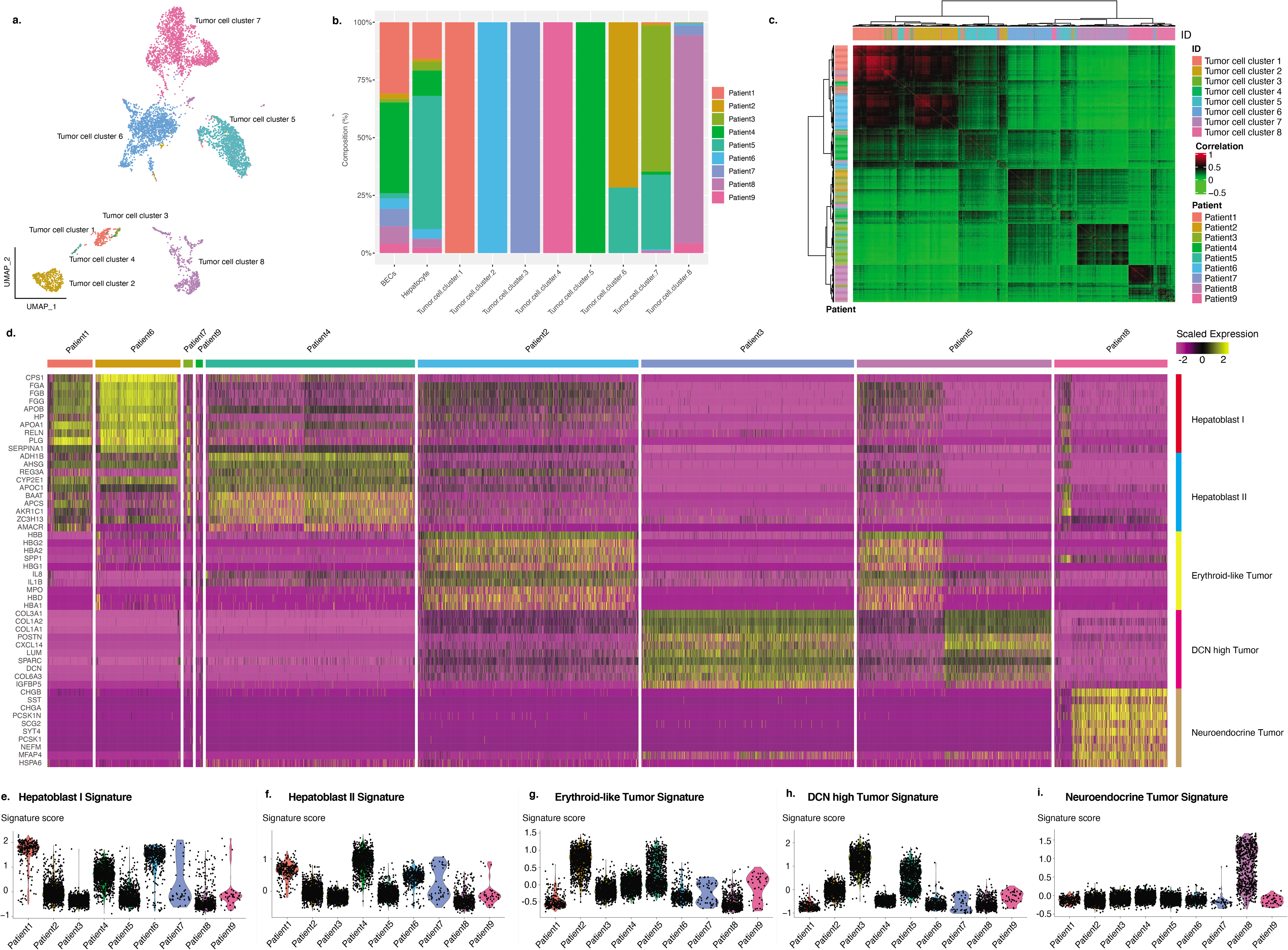
Tumor cell analysis revealed five transcriptomically distinct tumor cell types detected within the nine HB patients. a. Graphically clustering of all tumor cells (N = 6,244) by transcriptomic profiles. b. Stacked bar charts showing the contribution of the nine patients to each tumor population. c. Correlation heatmap of the tumor cells by tumor cell clusters and patients. Correlation was shown by the color gradient. Hierarchal clusters were illustrated by dendrograms. d. Heatmap of top 10 most differentially expressed genes of each tumor subtype in the nine patients. Tumor subtypes are annotated on the right. Scaled expressed levels are shown by the color bar. e. Violin plot of the computed Hepatoblast I tumor-subtype signature scores for all nine patients. f. Violin plot of the computed Hepatoblast II tumor-subtype signature scores for all nine patients. g. Violin plot of the computed erythroid-like tumor-subtype signature scores for all nine patients. h. Violin plot of the computed DCN-high tumor-subtype signature scores for all nine patients. i. Violin plot of the computed neuroendocrine tumor-subtype signature scores for all nine patients.

We investigated the similarity among tumor cells using a correlation heatmap with unsupervised clustering (**Figure 2c**), demonstrating that the tumor cells fell into five distinct groups. The epithelial tumor cell populations 1-4 were similar to each other and formed a distinct group. Group 2 consisted primarily of cells from the only pure fetal HB in our dataset. The three remaining groups consisted primarily of cells from the erythroid, *DCN*-high and neuroendocrine cell populations, respectively. Based on this, we generated five distinct HB tumor cell signatures (**Figure 2d, Supplemental Table 4**). Each of these five groups contained cells from multiple tumor cell populations, and were composed of cells from multiple patients, suggesting they identify common HB tumor cells shared across patients (**Supplemental Figure 6, Figure 2e-i**).

To further characterize these five HB tumor cell signatures, we compared them to published gene signatures obtained from bulk tissue gene expression datasets[8, 9]. We found that Hepatoblast I signature was enriched in both fetal and adult liver genes while Hepatoblast II signature was enriched only in adult liver genes (**Supplemental Figure 8**, p < 2.2e-16, two-sided Wilcoxon rank sum test). The Erythroid-like, *DCN*-high and Neuroendocrine signatures did not match signatures identified in previous studies. Gene ontology analysis further demonstrated differences between these five signatures by showing that Hepatoblast I and II signatures were enriched for metabolic processes typically found in hepatocytes, the Erythroid signature was enriched for immune and detoxification processes, the *DCN*-high signature was enriched for bone and extracellular matrix processes, and the Neuroendocrine signature was enriched for neuronal processes (**Supplemental Figure 8**).

Next, we asked if the five tumor cell signatures could explain the heterogeneity within each tumor. First, we characterized the degree of heterogeneity within each tumor (**Supplemental Figure 9**) and found that HBs have a range of heterogeneity within each tumor consistent with the observed pathology. Patient 4 tumor cells were the most homogeneous, consistent with its pure fetal histology, whereas Patient 8 tumor cells demonstrated the greatest degree of heterogeneity, consistent with its mixed epithelial mesenchymal histology. We generated a heatmap and showed that the five signatures corresponded well to most of the patient-specific tumor clusters (**Supplemental Figure 9**). This suggests that intratumor heterogeneity can be accounted for by the five tumor cell groups. We also detected subclusters that did not show an enrichment in any of the five tumor cell signatures, raising the possibility of other tumor subtypes (**Supplemental Figure 9**).

Finally, we asked whether the five tumor cell signatures can help predict tumor aggressiveness. We first used previously published predictive HB gene signatures and performed a pseudo-bulk RNAseq analysis of our nine tumor samples, and found that none of our 9 pseudo-bulk tumor transcriptomes were significantly enriched for either the low-risk rC1 or the high-risk rC2 gene sets (**Supplemental Figure 8**)[8]. Next, we repeated the analysis but used the five tumor signatures and found that Hepatoblast I and II showed significant enrichment for the low-risk rC1 gene set (p < 2.2e-16, two-sided Wilcoxon rank sum test), and the Neuroendocrine tumor subtype was enriched in the high-risk rC2 gene set (p = 5.37e-3, two-sided Wilcoxon rank sum test, **Supplemental Figure 8**). Interestingly, Patient 8, who contributed all HB tumor cells harboring the neuroendocrine signature, developed early relapse of HB within 12 months of surgical resection (Supplemental Table 1). This suggests that the five HB tumor subtype signatures may allow for more accurate risk stratification than bulk tumor transcriptomes.

### Low *MARCO* expression distinguishes HB tumor-associated macrophages from macrophages in normal tissue

We questioned whether differences in the tumor microenvironment were associated with the heterogeneity observed among tumor cells. We examined the immune and stromal cells in our dataset that were enriched within HB tumors (**Figure 3a,b**). Three immune cell populations were enriched within tumor, including macrophages, pro-myelocytes, and basophils (**Figure 3c**). Subset analysis of each of these three populations showed that tumor-associated macrophages (TAM) were the only cell type that exhibited significant transcriptomic differences compared to non-tumor macrophages (**Figure 3d, Supplemental Figure 10**). We compared the differentially expressed genes between macrophages from HB tumor tissue and adjacent non-tumor liver, and found that expression of the scavenger receptor, *MARCO*, distinguished macrophages found in tumors (*MARCO*^low^) from those isolated from adjacent normal liver (*MARCO*^Hi^) (**Figure 3e,f**). Notably, all nine patients contributed to the *MARCO*^low^ cluster in HB tumors (**Figure 3g**). We identified several differentially expressed genes that were upregulated in tumor-associated *MARCO*^low^ macrophages including the pro-tumorigenic chemokine *CCL18*[17] and genes known to promote cell proliferation, invasion, and migration (glycoprotein *NMB, GPNMB*[18], and FOS-related antigen 2, *FOSL2*[19](**Figure 3h,i**). Gene ontogeny pathway analysis demonstrated that *MARCO*^low^ TAMs had higher expression of genes involved with antigen presentation, chemotaxis, and processes related to protein production (**Figure 3j**).

**Figure 3.**
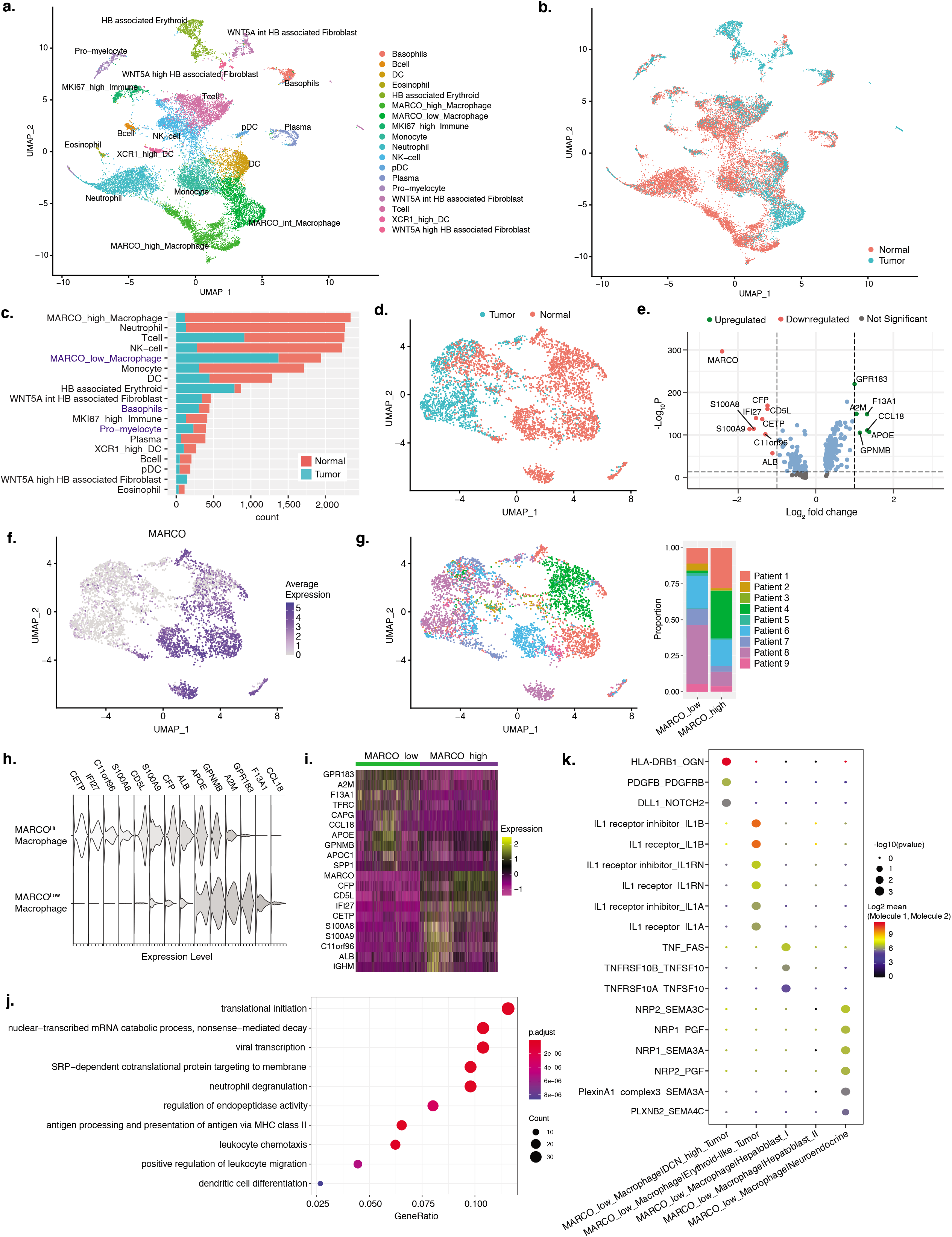
Low MARCO expression distinguishes HB TAMs from macrophages in normal tissue. UMAP of all immune cells and tumor-associated stromal cells from all 9 patients annotated by a. cell type or b. sample type. c. The quantities of each cell type in both tumor and adjacent normal tissue. d. UMAP of all macrophages from all 9 patients annotated by sample type. e. Volcano plot showing top differentially expressed genes between macrophages found in tumors compared to those found in adjacent normal liver tissue. f. Expression of the scavenger receptor, MARCO, distinguishes TAMs from normal liver macrophages. g. UMAP of all macrophages from all 9 patients annotated by patient (left) and the proportional contribution from each patient to the MARCO^Low^ and MARCO^Hi^ macrophage clusters. h. Violin plots showing expression of the genes identified in the above volcano plot in MARCO^Low^ TAMs and normal liver MARCO^Hi^ macrophages. i. Heatmap showing differentially expressed genes between MARCO^Low^ macrophages and MARCO^Hi^ macrophages. j. Gene ontology terms that are associated with the genes differentially expressed in MARCO^Low^ macrophages. k. Dot plot of significant ligand-receptor interactions between MARCO^Low^ macrophages and tumor epithelial cells for each patient. Mean expressions of the two molecules in the ligand-receptor pairs are depicted with color gradients and statistical significance is indicated by the dot sizes.

We next asked whether *MARCO*^low^ TAMs interact differently with each HB tumor cell subset identified in our analysis. We identified distinct ligand-receptor interactions between *MARCO*^low^ TAMs and four of the five HB tumor cell types. These distinct *MARCO*^low^ TAM interactions include PDGF signaling with *DCN*-high tumor cells, IL1 signaling with erythroid-like tumor cells, members of the TNF receptor superfamily in the Hepatoblast I tumor cell type, and plexin proteins (PLXNB2) with neuroendocrine-like tumor cells (**Figure 3k**). Collectively, these findings demonstrate that *MARCO*^Low^ TAMs express several pro-tumorigenic genes and that their unique interactions with the tumor cell types may contribute to the tumor heterogeneity observed in HB.

### Developmentally restricted erythroid progenitor cells associated with hepatoblastoma

The identification of an HB-associated erythroid population suggests that in postnatal livers, the tumor microenvironment of HB may maintain a fetal liver-like niche that results in persistence of erythroid progenitor cells. In our dataset, the HB-associated erythroid population was seen in three patients (Patients 2, 5 and 6). We first asked if HB-associated erythroid cells from these three patients had similar transcriptomes as human fetal liver erythroid cells. We integrated HB-associated erythroid cells with fetal liver erythroid cells from a reference dataset[14]. HB-associated erythroid cells shared gene expression profiles with human fetal liver erythroid cells and expressed markers from all three developmental stages, indicating that early, mid and late erythroblasts were present in the HB-associated population (**Figure 4a, b**).

**Figure 4.**
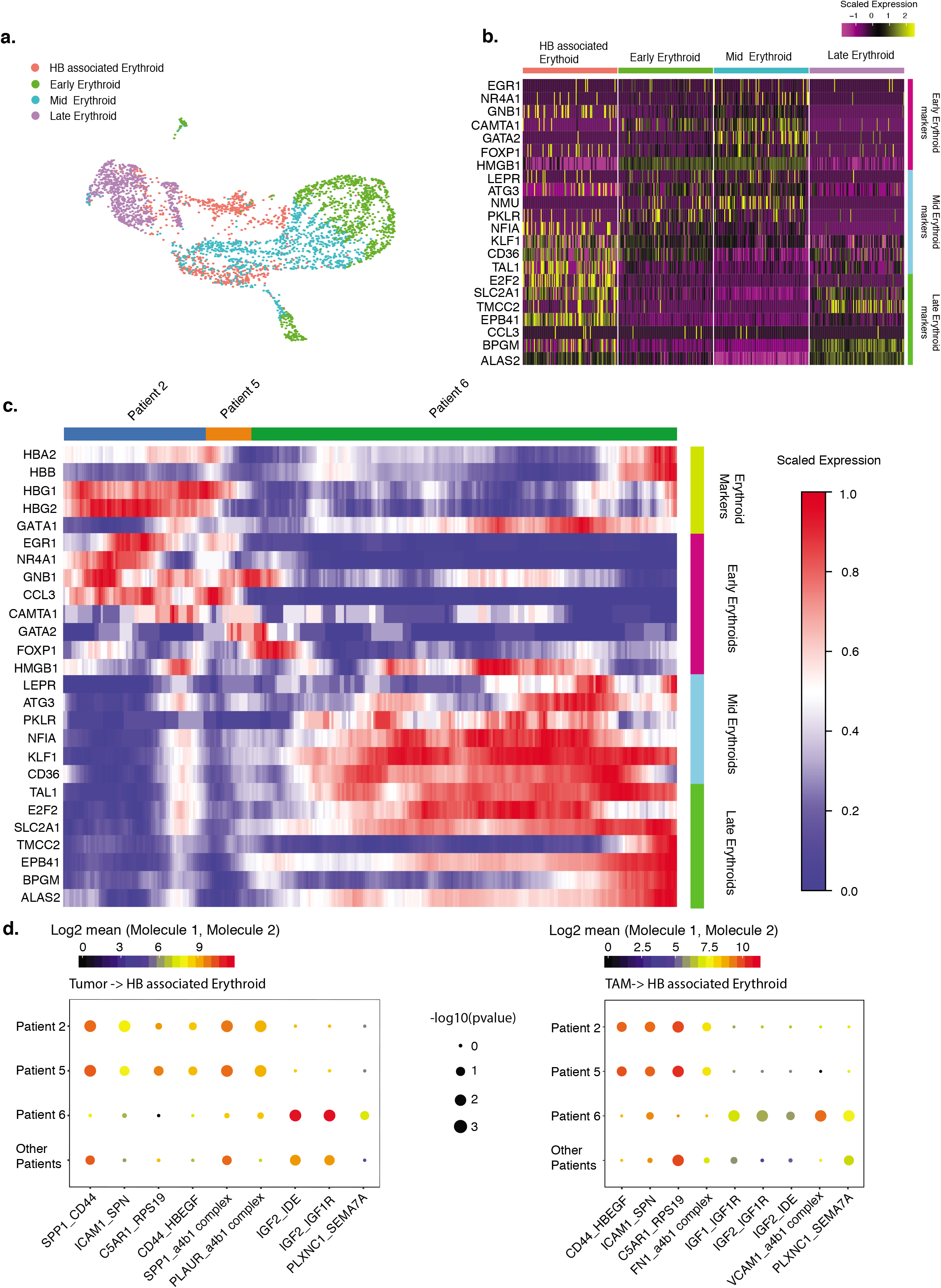
Analysis of erythroid and erythroid-like tumor cells maintained three fetal liver erythroid stages. a. Left panel, integrated UMAP of fetal liver erythroid cells from three developmental stages and HB-associated erythroid cells. Right panel, UMAP tumor cells and HB-associated erythroid cells from Patients 2, 5 and 6. b. Heatmap of the three developmental stages of fetal-liver erythroid marker expression for HB-associated erythroid cells and reference fetal liver erythroid population. Scaled expressed levels are shown by the color bar. c. Supervised heatmap showing expression of canonical erythroid markers and representative markers of three erythroid developmental stages in Patients 2, 5 and 6 HB associated erythroid cells. e. Left panel, representative ligand-receptor interactions between tumor cells and HB-associated erythroid cells for Patients 2, 5, 6, and other patients. Right panel, representative ligand-receptor interactions from TAMs to HB-associated erythroid cells for Patients 2, 5, 6, and other patients. Mean expressions of ligand and receptor pairs are shown in the color bar. Statistical significance levels are indicated by the marker size.

Next, we projected markers of fetal liver erythroid developmental stages with fetal erythroid markers *HBA2, HBB, HBG1* and *HBG2* on HB-associated erythroid cells from Patients 2, 5 and 6. We discovered that only cells from Patients 2 and 5 expressed early erythroid markers whereas cells from Patient 6 expressed mid and late erythroid genes (**Figure 4c**). We confirmed this by analyzing HB-associated erythroid cells separately and showed that cells from Patients 2 and 5 were predominantly early erythroids, whereas those from Patient 6 were primarily mid to late stage erythroid cells (**Supplemental Figure 11**).

We examined whether erythropoiesis in HBs progresses normally. Pseudotime analysis comparison between the reference fetal erythroid cells (**Supplemental Figure 11**) and HB-associated erythroid cells showed that cells from Patients 2 and 5 were arrested at the early erythroblast stage[20]. Cells from Patient 6, on the other hand, were at mid and late erythroblast stages.

Erythropoiesis in the fetal liver occurs within specialized niches composed in part by erythroblastic island macrophages and depends on intercellular interactions between the niche and erythroid progenitor cells that differ based on the specific stage of erythropoiesis[14, 21]. We hypothesized that TAMs and erythroid-like tumor cells from Patients 2, 5 and 6 may serve as erythroid niche cells. Ligand-receptor interactions (CellPhoneDB) between these two potential niche cell populations and the corresponding HB-associated erythroid cells demonstrated that TAMs and tumor cells from Patients 2 and 5 only expressed early erythroid niche signals, while those from Patient 6 only expressed late erythroid niche signals (**Figure 4d**)[22]. These results suggest that TAMs and erythroid-like tumor cells from these three patients form a niche and maintain HB-associated erythroid cells at their respective developmental stage. Overall, our data shows that HBs maintain fetal erythropoiesis at different developmental stages and supports the hypothesis that HB tumor heterogeneity is due in part to the fetal stage when the tumor arises[8, 11, 23].

We examined whether the tumor cells retained transcriptomic signatures corresponding to specific fetal liver developmental time points. We integrated tumor cells with a reference fetal liver hepatoblast dataset but found little overlap, indicating that tumor cells had gene expression patterns distinct to hepatoblasts (**Supplemental Figure 12)**[16, 24].

We noticed that Patients 2 and 5 had tumor cells with expression of erythroid genes but Patient 6 did not. We ensured that the presence of erythroid genes in these cells was not due to ambient RNA contamination (**Supplemental Figure 13**) or doublets (data not shown). Erythroid-like tumor cells in Patients 2 and 5 expressed early erythroblast markers, but had lower expression of more mature erythroid markers *HBG1, HBG2, HBA2, and HBB* than HB-associated erythroid cells. Tumor cells from Patient 6 showed little to no expression of most erythroid markers, consistent with our previous analysis that Patient 6 tumor cells were predominantly hepatoblast-like (**Supplemental Figure 13**). Finally, we examined ligand-receptor interactions between TAMs and tumor cells and found erythropoiesis signals were only present between TAMs and tumor cells for Patients 2 and 5 (**Supplemental Figure 13**). This suggests that erythropoietic signaling may be involved in the oncogenesis for at least a subset of HBs.

### Patient-derived-spheroids maintain patient-specific features from freshly isolated parent cells

We used a recently developed protocol to generate five patient-derived spheroids (PDS) (Peng Wu, Roel Nusse unpublished data). The success rate was greater when PDS were grown from freshly isolated cells than from cryopreserved cells (**Supplemental Table 5**). Four of the five PDS were maintained for more than 15 passages (>6 months). The PDS from Patient 9 had two distinct cell populations corresponding to non-tumor BECs and fibroblasts, neither of which could be maintained long-term in our culture conditions (**Supplemental Figure 14**). PDS from different patients had distinct morphologies, which persisted with long-term culture (**Figure 5a**, **Supplemental Figure 14**). PDS had similar proliferation rates which remained stable over time (**Supplemental Figure 14**).

**Figure 5.**
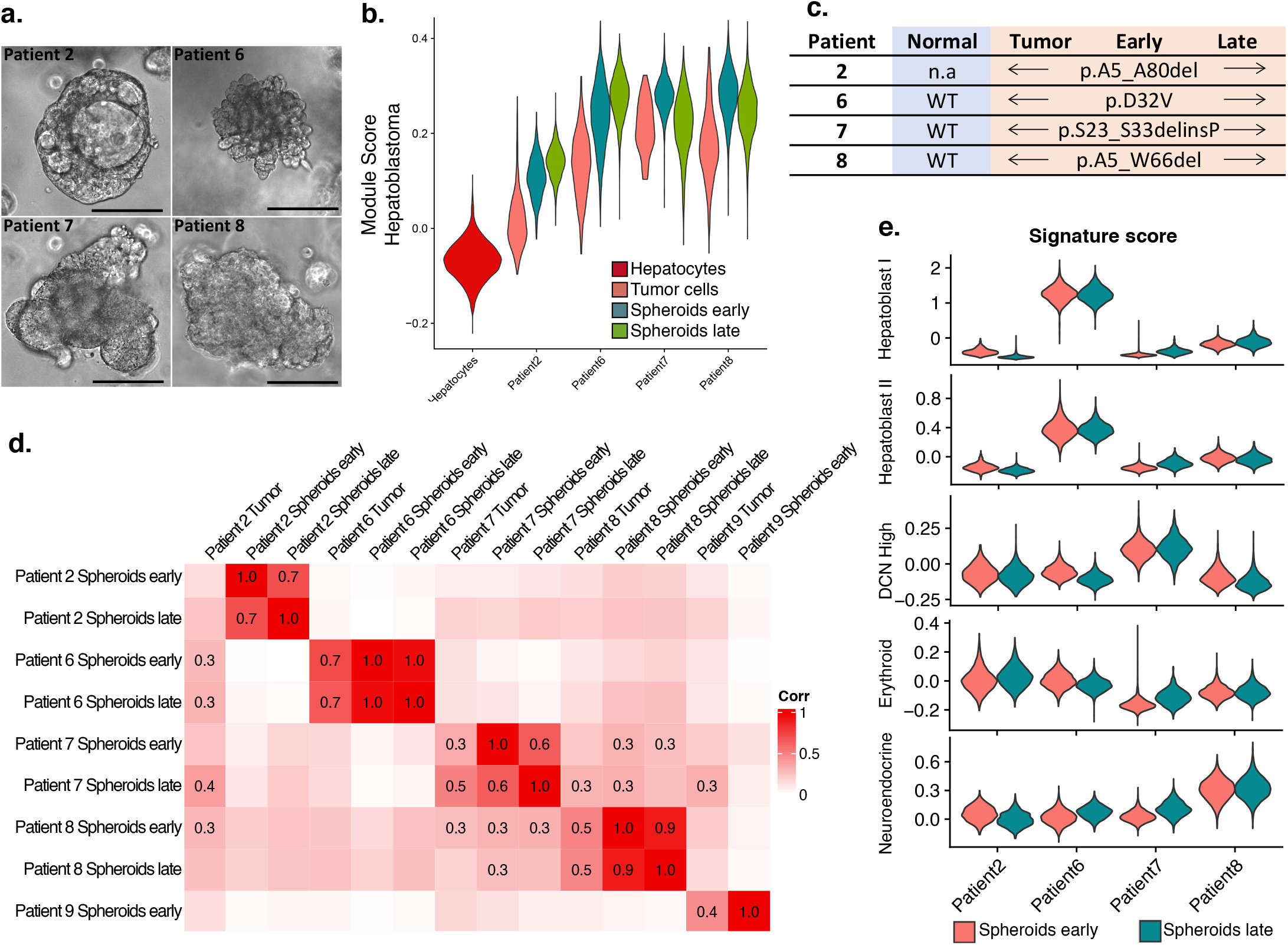
Patient-derived-spheroids maintain patient-specific features from freshly isolated parent cells. a. Brightfield images of tumor spheroids from Patients 2, 6, 7 and 8. Scale bar = 100 μm. b. Violin plots of KEGG Cairo_Hepatoblastoma_UP signature score of tumor cells (Patient 2) or spheroid parent cells (Patient 6, 7 and 8) or spheroids from Patients 2, 6, 7 and 8. Signature scores for all hepatocytes from all patients are represented as a reference. c. Exon 3 somatic mutation of *CTNNB1* (#NP_001091679.1) of normal and tumor tissues, spheroids, early and late passages. d. Correlation matrix between PDS and freshly isolated tumor cells. e. PDS module scores (early and late passages) for the signatures of the five tumor cell types.

To further characterize the PDS, we performed scRNA-seq on spheroids at both early and late passages. PDS have broad gene expression differences compared to freshly isolated tumor cells, but retained their transcriptomic differences relative to other PDS even with long-term culture, indicating that they maintained patient-specific features (**Supplemental Figure 15**). All PDS had high expression of hepatoblastoma genes and retained their parent tumor mutations in *CTNNB1* (**Figure 5b,c**), indicating that they grew from tumor cells. To directly assess which specific cell population each PDS grew from, we generated a signature score for each cell population in the tumor tissues and found that PDS were most similar to the tumor cell population in each case, even after long-term culture (**Figure 5d, Supplemental Figure 14**). We could not identify a clear cell of origin for PDS from Patient 2, likely because we did not capture the full spectrum of tumor cells in this patient (**Supplemental Figure 7**).

PDS from Patient 7 had two distinct cell populations, a large population of spheroid cells similar to tumor cells, and a small population similar to fibroblasts. This is consistent with the finding of fibroblast-appearing cells in early passages for Patient 7, which disappeared with long-term culture (**Supplemental Figure 14**).

We next asked if PDS retained gene expression of any of the five HB tumor cell types we identified. Since we did not capture the parent cells for PDS from Patient 2 and very few tumor cells were captured from Patient 7, we focused our analysis on Patients 6 and 8. We analyzed the tumor cells from Patients 6 and 8 and generated a signature score for each subpopulation (**Supplemental Figure 16**). PDS from Patient 6 is most similar to the population of tumor cells that expressed the Hepatoblast I tumor cell signature. Patient 8 PDS is most similar to the population of tumor cells with Neuroendocrine tumor cell signature. This suggests that each PDS grows from a specific subpopulation of tumor cells, which is consistent with the relative homogeneous transcriptome profile of PDS. We measured the gene signature score for each PDS relative to the five HB tumor cell signatures and showed that PDS from Patients 6, 7 and 8 likely grew from the five HB tumor cell types we identified, but Patient 2 did not (**Figure 5e**).

### Pharmacologic testing of patient-derived HB spheroids reveals HB tumor cell type specific treatment responses

We examined whether PDS can be used to test drug-sensitivity in a patient-specific manner. We treated PDS with five chemotherapeutics commonly used for HB and found that spheroids from Patient 7 were the most sensitive to the drugs tested, while PDS from Patients 2 were the most resistant (**Figure 6a**). Drug sensitivity is partly dependent on the cell’s ability to metabolize and efflux the agent. We measured expression levels of genes known to be important for efflux and metabolism of platinum-based drugs, etoposide, vincristine, and 5-FU (https://www.pharmgkb.org/pathway/PA150653776, https://www.pharmgkb.org/pathway/PA150981002, https://www.pharmgkb.org/pathway/PA2025, https://www.pharmgkb.org/pathway/PA150981002) and found that in both tumor cells and spheroids, Patient 2 had the highest expression of these genes while Patient 7 had the lowest (**Figure 6d, Supplemental Figure 17**).

**Figure 6.**
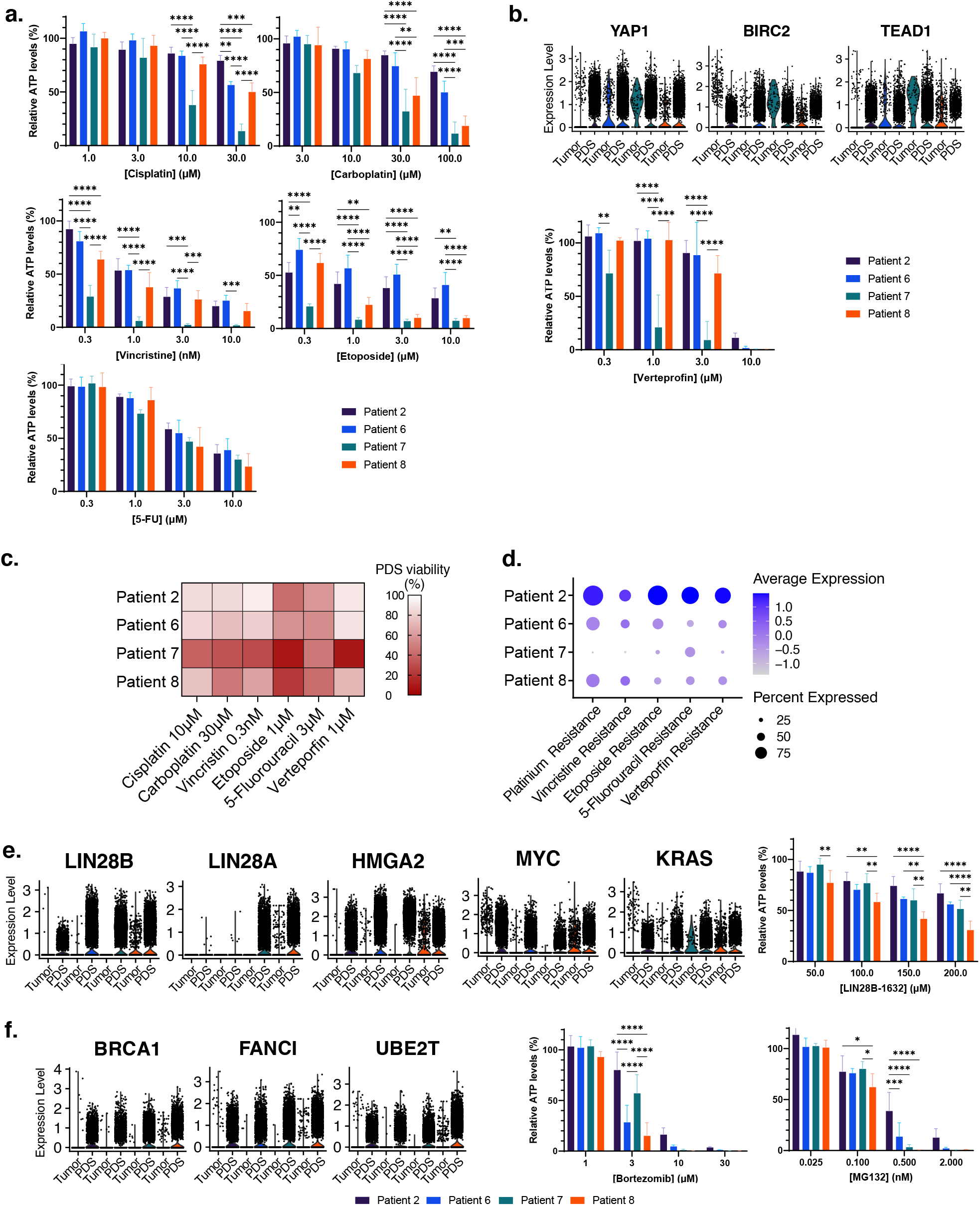
Pharmacologic testing of patient-derived HB spheroids reveals HB tumor cell-type-specific treatment responses. a. Cell viability measured by ATP quantification after a 4-day treatment with five chemotherapy drugs (cisplatin, carboplatin, etoposide, vincristine and 5-fluorouracil [5-FU]) b. Violin plots showing the expression of YAP1, BIRC2 and TEAD1 in-parent cells and PDS, and cell viability measured by ATP quantification after a 4-day treatment with the YAP1 inhibitor verteporfin. c. PDS drug-sensitivity heatmap showing cell viability profile for the five chemotherapy drugs tested and verteporfin at a representative concentration. d. Module score for genes involved in platinum-based compounds, etoposide, vincristine, 5-FU, and verteporfin metabolism and efflux. e. Violin plots showing the expression of LIN28B, LIN28A, HMGA2, MYC and KRAS in-parent cells and PDS, and cell viability measured by ATP quantification after a 4-day treatment with the LIN28 pathway inhibitor LIN28B-1632. f. Violin plots showing the expression of BRCA1, FANCI and UBE2T in-parent cells and PDS, and cell viability measured by ATP quantification after a 4-day treatment with the BRCA-FA pathway inhibitors bortezomib and MG132.

Next we asked whether PDS can be used to identify novel treatment targets for HB. We looked for known molecular pathways important for HB pathogenesis and are differentially expressed within our HB tumors and corresponding PDS. We found that the YAP signaling pathway was highly expressed in Patient 2 and 7 cells (**Figure 6b**). We treated PDS with the YAP1 inhibitor verteporfin and found that Patient 7 PDS were most responsive to YAP1 inhibition whereas Patient 2 PDS were not (**Figure 6b**). To explain this discordance, we looked at the expression of genes known to be important for efflux and metabolism of verteporfin and found that Patient 2 had the highest expression of these genes (**Figure 6d, Supplemental Figure 17**)[25]. Taken together, these results allowed us to generate a drug sensitivity profile of PDS that is globally linked to the expression of drug efflux and metabolism genes in the tumor cell and PDS (**Figure 6c, d**).

Since Patient 8 had an early relapse, we looked specifically at signaling pathways differentially expressed in Patient 8 tumor and PDS. We found Patient 8 cells expressed genes in LIN28B and BRCA/FANC pathway. We treated the PDS with inhibitors to both pathways and found that Patient 8 cells were sensitive to both (**Figure 6e, f**).

## DISCUSSION

Hepatoblastoma is a childhood cancer in which the response to chemotherapy can be limited by high-levels of chemotherapeutic resistance[26]. We postulate that an important determinant of the variability in chemo-sensitivity is the cellular composition of individual tumors, and have utilized the power of single cell genomics to i) define the heterogeneity of pediatric HB and ii) relate it to chemotherapeutic efficacy. We identified five transcriptomically distinct tumor cell signatures that are present in varying proportions across all tumors, indicating that the relative abundance of these tumor cells underlies the heterogeneity observed in HB. We also identified specific tumor-associated cells, including TAMs, that form the tumor microenvironment. In a subset of HBs, TAMs and tumor cells form a unique niche to maintain fetal liver erythropoiesis. Interestingly, TAMs and tumors express developmentally restricted signals that result in similarly restricted erythropoiesis. Finally, we grew patient-derived spheroids from HB tumors and found that each spheroid culture was an outgrowth of one of the five tumor cell types, establishing a useful tool to study HB biology and drug susceptibility. Taken together, this study identifies the tumor cells types that drive heterogeneity in this disease and establishes tools to improve our understanding of HB biology and its treatment.

The clinical outcomes for HB depends on the presence of specific epithelial cell types, such as for those patients with small cell undifferentiated histology[27, 28]. Consistent with this, bulk transcriptome data separate tumor risk primarily by epithelial component gene expression[8]. Our approach identified two epithelial tumor cell types shared across HB, including cells with the pure fetal histology. We also identified three new tumor cell types corresponding to cells in mixed epithelial mesenchymal tumors. This includes a rare neuroendocrine population which correlated with high clinical risk in our dataset. Using this approach on a larger HB dataset could identify most if not all HB tumor cell types. By correlating these with clinical outcomes, it will be possible to define gene signatures that accurately risk stratify HB.

HBs are thought to arise from fetal hepatoblasts, and are known to have few genetic mutations, most in the Wnt signaling pathway, suggesting that tumor heterogeneity in HB is not driven by distinct mutations[29]. In this study, we demonstrate that HBs maintain features that correspond to specific fetal liver developmental stages. This finding supports the hypothesis that HB heterogeneity is determined by the stage at which the tumor arose during liver development[8, 11, 23].

Patient-derived organoids and spheroids are powerful models to study tumor biology[30, 31]. We showed that PDS can be grown efficiently from fresh HB tumor samples and retain patient and tumor cell type gene expression patterns even after long-term culture. Importantly, PDS predict differential responses to treatment based on the transcriptomic signature of each tumor, suggesting a path forward for precision oncology for these tumors.

Our data is limited by the small sample size of nine tumors, and the fact that eight of our samples are post-treatment, which likely affected the cell types captured and perhaps enriches for chemo-resistant tumor cell populations. In addition, it is likely that additional HB tumor cell types exist which we were not able to capture, such as the tumor cells that gave rise to PDS from Patient 2. Recent developments in single nuclei RNA sequencing may allow for examination of larger biobanks of HB samples, which we expect will lead to the identification of other common HB cell types[32].

In conclusion, our results define HB tumor heterogeneity with single-cell resolution and demonstrate that patient-derived spheroids can be used to evaluate responses to chemotherapy.

## ONLINE METHODS

### Human specimen collection

Fresh tumor tissue and adjacent normal tissue from nine patients were obtained during anatomic liver resections for hepatoblastoma. Tissues where then kept during transportation in ice-cold Williams’ E medium (ThermoFisher Scientific, Waltham, MA) (supplemented with 2 mM Glutamax, 10 mM HEPES and 1000U/ml Penicillin/Streptomycin (ThermoFisher Scientific)), and then processed for scRNA-seq and spheroid culture. This study was approved by the UCSF Institutional Review Board (IRB).

### Single-cell RNA sequencing

Tumor and adjacent normal liver were dissociated by mincing tissue in 1-2mm squares, followed by incubation in Liver Perfusion Medium (ThermoFisher Scientific) for 15 min at 37°C in a rotating oven. After being washed in PBS (ThermoFisher Scientific), tissues were incubated with Liver Digest Medium (ThermoFisher Scientific) supplemented with HEPES and collagenase type IV (Worthington Biochemical, Lakewood, NJ) (600 to 800 U/ml) for 30 min at 37°C with rotation. Further dissociation was achieved by pipetting 10 times through a 25 mL serological pipette. Single cells were then separated from clumps using a 70 μm strainer (Fisher Scientific, Hampton, NH). After lysis of red blood cells with ACK RBC Lysis Buffer (Fisher Scientific), single cells were counted using a LUNA™ Automated Cell Counter (Logos biosystems, South Korea) and processed for scRNA-seq analysis.

Sequencing was based on the Seq-Well S^3 protocol [33]. One to four arrays were used per sample. Each array was loaded as previously described with approximately 110,000 barcoded mRNA capture beads (ChemGenes, Cat: MACOSKO-2011-10(V+)) and with 10,000-20,000 cells. Arrays were then sealed with functionalized polycarbonate membranes (Sterlitech, Cat: PCT00162X22100) and were incubated at 37°C for 40 minutes in lysis buffer (5 M Guanidine Thiocyanate, 1 mM EDTA, 0.5% Sarkosyl, 1% BME). After detachment and removal of the top slides, arrays were rotated at 50 rpm for 20 minutes. Each array was washed with hybridization buffer (2 M NaCl, 4% PEG8000) and then incubated in hybridization buffer for 40 minutes. Beads from different arrays were collected separately. Each array was washed ten times with wash buffer (2 M NaCl, 3 mM MgCl_2_, 20 mM Tris-HCl pH 8.0, 4% PEG8000) and scraped ten times with a glass slide to collect beads into a conical tube.

For each array, beads were washed with Maxima RT buffer (ThermoFisher, Cat: EP0753) and resuspended in a master mix comprised of Maxima RT buffer, PEG8000, Template Switch Oligo, dNTPs (NEB, Cat: N0447L), RNase inhibitor (Life Technologies, Cat: AM2696) and Maxima H Minus Reverse Transcriptase (ThermoFisher, Cat: EP0753) in water. Samples were rotated first at room temperature for 15 minutes and then at 52°C overnight. Beads were washed once with TE-SDS and twice with TE-TW. They were treated with exonuclease I (NEB), rotating for 50 minutes at 37°C. Beads were then washed once with TE-SDS, twice with TE-TW, and once with 10 mM Tris-HCl pH 8.0. They were resuspended in 0.1 M NaOH and rotated for five minutes at room temperature. They were subsequently washed with TE-TW and TE, and taken through second strand synthesis with Maxima RT buffer, PEG8000, dNTPs, dN-SMRT oligo and Klenow Exo- (NEB, Cat: M0212L) in water. After rotating at 37°C for 1 hour, beads were washed twice with TE-TW, once with TE and once with water.

KAPA HiFi Hotstart Readymix PCR Kit (Kapa Biosystems, Cat: KK2602) and SMART PCR Primer were used in whole transcriptome amplification (WTA). For each array, beads were distributed among 24 PCR reactions. Following WTA, three pools of eight reactions were made and were then purified using SPRI beads (Beckman Coulter), first at 0.6x and then at a 0.8x volumetric ratio. For each sample, one pool was run on an HSD5000 tape (Agilent, Cat: 5067-5592). The concentration of DNA for each of the three pools was measured via the Qubit dsDNA HS Assay kit (ThermoFisher, Cat: Q33230). Libraries were prepared for each pool, using 800-1000 pg of DNA and the Nextera XT DNA Library Preparation Kit. They were dual-indexed with N700 and N500 oligonucleotides.

Library products were purified using SPRI beads, first at 0.6x and then at a 1x volumetric ratio. Libraries were then run on an HSD1000 tape (Agilent, Cat: 50675584) to determine the concentration between 100-1000 bp. For each library, 3 nM dilutions were prepared. These dilutions were pooled for sequencing on a NovaSeq S4 flow cell.

The sequenced data were preprocessed and aligned using the dropseq workflow on Terra (app.terra.bio). A digital gene expression matrix was generated for each sample, parsed and analyzed following a customized pipeline.

### Sequencing and alignment

Sequencing results were returned as paired FASTQ reads and the paired FASTQ files were aligned against hg19 reference using the dropseq workflow (https://cumulus.readthedocs.io/en/latest/drop_seq.html). The aligning pipeline output included aligned and corrected bam files, two digital gene expression (DGE) matrix text files (a raw read count matrix and a UMI-collapsed read count matrix where multiple reads that matched the same UMI would be collapsed into one single UMI count) and text-file reports of basic sample qualities such as the number of beads used in the sequencing run, total number of reads, alignment logs. For each sample, the median and average number of genes per barcode were 1,014 and 1,224. The median and average number of UMI were 2,536 and 3,974. The mean percentage of mitochondrial content per cell was 17.06%.

### Single-cell clustering analysis

Cells captured in single-cell RNA sequencing analysis were clustered and analyzed using customized codes based on the Seurat (Version 3.2) package in R (Version 4.0.3) [34]. Cells with fewer than 300 genes, 500 transcripts, or a mitochondrial level of 20% or greater, were filtered out as the first QC process. Then, by examining the distribution histogram of the number of genes per cell in each sample, we set the upper threshold for the number of genes per cell in each individual sample in order to filter potential doublets. A total of 29,968 cells were acquired using these thresholds.

UMI-collapsed read-count matrices for each cell were loaded in Seurat for analysis. We followed a standard workflow by using the “LogNormalize” method that normalized the gene expression for each cell by the total expression, multiplying by a scale factor 10,000. For downstream analysis to identify different cell types, we then calculated and returned the top 2,000 most variably expressed genes among the cells before applying a linear scaling by shifting the expression of each gene in the dataset so that the mean expression across cells was 0 and the variance was 1. This way, the gene expression level could be comparable among different cells and genes. Principal components analysis (PCA) was run using the previously determined most variably expressed genes for linear dimensional reduction and the first 100 principal components (PCs) were stored, which accounted for 40.49% of the total variance. To determine how many PCs to use for the clustering, a JackStraw resampling method was implemented by permutation on a subset of data (1% by default) and rerunning PCA for a total of 100 replications to select the statistically significant PC to include for the K-nearest neighbors clustering. For graph-based clustering, the first 75 PC and a resolution of 1.2 were selected, yielding 37 cell clusters. We eliminated the clustering side effect due to over clustering by constructing a cluster tree of the average expression profile in each cluster and merging clusters together based on their positions in the cluster tree. As a result, we ensured that each cluster would have at least 10 unique differentially expressed genes (DEGs). Differentially expressed genes in each cluster were identified using the FindAllMarker function within Seurat package and a corresponding p-value was given by the Wilcoxon’s test followed by a Bonferroni correction.

### Cell type signature analysis

In order to annotate each cell type from the previous clustering, we referred to established studies and used signature gene sets for each cell type (**Suppl Table 2**). Treating the signature gene set for each cell type as a pseudogene, we evaluated the signature score for each cell in our dataset using the AddModuleScore function. Each cluster in our dataset was assigned with an annotation of its cell type by top signature scores within the cluster. To validate the identities of the tumor cell populations, we estimated copy number variants (CNV) via InferCNV (Version 1.4.0), using non-tumor and non-tumor-associated populations as reference [15]. During the inferCNV run, genes expressed in fewer than five cells were filtered from the data set and the cut off was fixed at 0.1. Hidden Markov model (HMM) based CNV prediction was achieved and estimated CNV events were shown in a heatmap.

All tumor cells were subset and re-clustered using the analytical workflow described above. Eight clusters of tumor cells were obtained with distinctive transcriptomic profiles. By downsampling each cluster to 200 or fewer cells, we computed the correlation matrix between each tumor cell pairs and used the pheatmap R package (Version 1.0.12) (Kolde, 2019, https://CRAN.R-project.org/package=pheatmap) to make the correlation heatmap with unsupervised clustering. Gene signatures were tested on all eight tumor cell clusters using previously established hepatoblast, fibroblast, erythroid and neuroendocrine signature gene sets. A one-way ANOVA test was conducted for each signature score comparison across all eight tumor cell clusters and a corresponding p-value was computed.

In this analysis, we created customized gene set signatures for each cell population of interest. Using the DEGs obtained from FindAllMarker function, we included genes with log2 fold change > 2 and statistical significance (FDR q < 0.05) as the signature gene set.

### Tumor-associated erythroid developmental analysis

Tumor-associated erythroid populations were extracted and integrated with the fetal liver erythroid or erythroblast populations from two publicly available datasets ([24]descartes.brotmanbaty.org, human fetal liver erythroblast) ([14] fetal liver early, mid and late erythroid, ArrayExpress, accession code E-MTAB-7407). We utilized the integration method based on commonly-expressed anchor genes by following the Seurat integration vignette to remove batch effects of samples sequenced with different technologies and possible artifacts so that the cells were comparable.

To evaluate the tumor-associated erythroid and tumor cell populations with respect to the fetal developmental stages, we first calculated a partition-based graph abstraction (PAGA) graph using SCANPY’s (Version 1.4.2) (Wolf et al., Genome Biology, 2018) sc.tl.paga() function and then used sc.tl.draw_graph() to generate the PAGA-initialized single-cell embedding of the cell types. The expression of markers was projected from the three fetal erythroid developmental markers (early, mid and late) to each tumor and erythroid cluster to generate a heatmap.

### Pseudotime analysis

The erythroid population was exported as a Seurat object and then converted into a SingleCellExperiment sim object. Pseudotime analysis was conducted using the slingshot R package (Version 1.6.1) [20]. First, PCA decomposition was performed using the prcomp() function in the stats R package (Version 3.6.2). A diffusion map was then generated using the top layer annotation from the original Seurat object and the pseudotime trajectory was superimposed on the diffusion map. The starting point of the pseudotime trajectory was determined as the early erythroid population.

### Cell-cell interaction analysis

We evaluated cell-cell interactions between two populations of interest using the CellPhoneDB package (Version 2.1.4) (Efremova et al., Nature, 2020). For each analysis, two input files were generated including a normalized gene expression matrix and a two-column metadata for cell names and annotations. The normalized gene expression matrix was obtained by using the NormalizeData function in Seurat with “RC” method specified. Statistical analysis of all available ligand-receptor pairs was performed on local computers.

To investigate the biologically relevant cell populations, we filtered the CellPhoneDB p-value.txt output file for ligand receptor pairs with the p-value less than 0.05, indicating statistically significant interactions, and generated customized columns and rows txt files. Dot plots were then plotted using these files to illustrate only the significant ligand-receptor interactions.

### Data and code availability

Processed single-cell RNA sequencing data that support this study will be deposited in the NCBI GEO database and available upon request to the corresponding author. All software algorithms used for analysis are available for download from public repositories. All code used to generate figures in the manuscript are available upon request.

### Patient-derived HB tumor spheroids

Patient-derived spheroids were cultivated either from cryopreserved single cells (Patient 2) or from fresh remaining clumps after tissue dissociation (Patients 6, 7, 8, 9). Cryopreserved cells were thawed in wash medium (Advanced DMEM/F12 Medium ((ThermoFisher Scientific) containing 2 mM Glutamax, 10 mM HEPES, 1,000 U/mL penicillin/streptomycin and 5% FBS (Fisher Scientific)). Then, cells were centrifugated at 400g for 5 min, and resuspended in tumor medium (Advanced DMEM/F12 supplemented with 2 mM Glutamax, 10 mM HEPES, 1,000U/mL penicillin/streptomycin, 2% B27 (ThermoFisher Scientific), 1% N2 (ThermoFisher Scientific), 10 mM Nicotinamide (MilliporeSigma, Burlington, MA), 1.25 mM N-acetylcysteine (MilliporeSigma), 10 μM Y27632 (BioGems, Westlake Village, CA), 100 ng/mL hFGF10 (PeproTech, Rocky Hill, NJ), 25 ng/mL hHGF (PeproTech), 50 ng/mL hEGF (PeproTech), 5 μM A83-01 (Fisher Scientific) and 3 nM dexamethasone (BioGems)). Cells were first seeded in a low binding plate for 4 hours at 37°C, 5% CO_2_, to promote cell clumping, then clumps were centrifugated, resuspended in pure Matrigel® and 25-μL domes were seeded in 48-well plates. After allowing the Matrigel® (Corning, Corning, NY) to solidify at 37°C, tumor medium was added. For fresh culture, tissue clumps were centrifugated and resuspended in pure Matrigel®, and seeded as previously described. Medium was renewed every 3 to 4 days. Spheroids were visible after 2 to 4 days and passaged after 2 weeks.

Subsequent passaging was performed every 7 to 10 days by splitting cells at a ratio of 1:10 to 1:30. Briefly, domes were washed with PBS, Matrigel® was digested, and cells were dissociated with TrypLE 1X (ThermoFisher Scientific) at 37°C for 20 to 45 minutes (varying with cell line) until spheroids became small clumps. Then, cells were centrifugated for 5 min at 400 g at 4°C and rinsed with wash medium. The cell pellet was then resuspended in pure Matrigel®, and seeded as previously described.

Expression profiling of spheroids was performed by scRNA-seq analysis at an early passage (2-3) and at a late passage (10-11). Briefly cells were prepared as for passaging, using an extended incubation in TrypLE 1X to obtain a single cell suspension. Then, cells were filtered through a 70-μm strainer, counted using a LUNA™ Automated Cell Counter, and loaded into a Seq-Well array as previously described. A total of 15,922 cells passed the QC step from all five cell lines, with an average nGene of 3,570 and an average nUMI of 10,264.

Doubling time and drug cytotoxicity were evaluated based on ATP quantification (CellTiter-Glo® 3D Cell Viability Assay kit, Promega, Madison, WI). Briefly, cells were seeded at 25,000 cells per dome. For doubling time calculation, ATP content was measured in domes at days 1, 3, 5, 7, and 9 after seeding. Doubling time was calculated during the exponential phase of proliferation, namely between day 1 and day 5. For cytotoxicity drug testing, cells were seeded as previously described and treated with drugs at 4 different concentrations for 4 days, between day 1 and day 5. For each separate experiment, ATP levels were quantified using an ATP-standard curve. Cisplatin and carboplatin were reconstituted in PBS and all other drugs (vincristine, etoposide, verteporfin and 5-fluorouracil, bortezomib, MG132 and LIN28B-1632) in DMSO (Cell Signaling Technology, Danvers, MA). Controls were incubated with their respective vehicles (PBS or DMSO). Data show average +/- SD of 5 independent experiments, and correspond to the percentage of ATP level normalized to the control.

#### CTNNB1 mutation detection

DNA was extracted from frozen tissues or cryopreserved freshly isolated cells for tumor and tumor-adjacent samples, and from spheroids at early and late passages, using DNA/RNA All Prep mini Kit (Qiagen, Hilden, Germany). *CTNNB1* genomic sequence (from exon 2 trough exon 4) was amplified using specific primers (Sumazin, Hepatology, 2016) (forward primer: AGCGTGGACAATGGCTACTCAA; reverse primer: ACCTGGTCCTCGTCATTTAGCAGT) by polymerase chain reaction using Q5® Hot Start High-Fidelity 2X Master Mix (NEB, Ipswich, Ma), then sequenced by Sanger sequencing (MCLAB, South San Francisco, CA) using the same primers.

## ACKNOWLEDGEMENTS

We thank Pam Derish, Xin Chen, and Monty Bissell for their critical review and helpful comments of the manuscript. This work was supported by the American Cancer Society Individual Research Grant (IRG-17-180-1, AN), American Association for the Study of Liver Diseases Clinical, Translational, and Outcomes Research Award (AN), NIH K08DK101603 (BMW), Burroughs Wellcome Fund Career Award for Medical Scientists (BMW), and core resources of the UCSF Liver Center (P30 DK026743).

## AUTHOR CONTRIBUTIONS

HS and SB equally contributed to experimental design, performing the experiments, data analysis and interpretation, and writing of the manuscript.

KR performed the experiments and contributed to data analysis and interpretation.

MT performed the experiments and contributed to data analysis and interpretation.

DB performed the experiments.

SJC contributed to data analysis and interpretation.

AR contributed to data analysis and interpretation.

FWH, AN, and BW equally contributed to experimental design, data analysis and interpretation, writing of the manuscript, and supervising the work.

## REFERENCES

<60 refs, additional 20 can go into online methods):

## Supplemental Data

**Supplemental Figure 1. Quality control of cells.** a. Stacked bar charts of cells by overall quality for all patients. Low-quality cells are removed from downstream analysis. b. Violin plots of UMI counts per cell by patient. c. Violin plots of gene counts per cell by patient. d. Violin plots of percentage mitochondrial per cell by patient.

**Supplemental Figure 2. Clustering of all cells passing QC from nine HB patients.** a. UMAP of all cells passing QC annotated by PCA-based clustering. b. Stacked bar chart of cell counts from tumor and normal samples for nine patients c. Heatmap of the top five most differentially expressed genes for each cluster. Scaled expression is shown in the color bar.

**Supplemental Figure 3. Clustering by sample types and patients from nine HB patients.** a. UMAP by sample types (tumor or adjacent normal samples) and stacked bar charts of each annotated clusters by sample types. b. UMAP by patients and stacked bar charts of each annotated cluster by patients.

**Supplemental Figure 4. Correlation heatmap of cell-type manual annotation and singleR automated annotation.** Heatmap showing the correlation coefficient between manual annotation and singleR automated annotation. SingleR annotations are shown on the left and manual annotation at the bottom.

**Supplemental Figure 5. Hepatoblastoma tumor gene expression pattern.** a. Dot plot of 8 hepatoblastoma genes in all cell populations. b. Dot plot of the top 10 most differentially expressed genes within the DCN-high tumor cell cluster and two fibroblast clusters. Average expression levels are shown in the color bar and percent of expression is indicated by the marker size. c. Violin plots of four representative hepatoblastoma tumor markers in the three clusters.

**Supplemental Figure 6. Tumor cell analysis.** a. UMAP of tumor cells by patients. b. stacked bar charts of the eight tumor cell clusters and the two non-malignant epithelial cell types by patients. c. Heatmap of top 10 most differentially expressed genes of the eight tumor clusters. Scaled expression levels are shown in the color bar. d. Feature plots of the five tumor subtype signature scores within the tumor cell UMAP. Computed signature scores are shown in the color bar.

**Supplemental Figure 7. Tumor tissue histology.** a-i: H&E stainings showing representative histological features of tumor tissues for patients 1 to 9. Magnification: 200x (a, e, f, h); 100x (b, c, d, g, i).

**Supplemental Figure 8. Gene ontology analysis of the five HB tumor subtypes and comparison with established HB signatures.** a. Box plots of Module 24, a human fetal liver gene set, and Hsiao liver-specific genes, a human adult liver gene set signature scores of tumor cells in the five tumor subtypes. b. Dot plots of the gene ontology analysis results for the five HB tumor subtypes. Top 20 ontology pathways are shown. Statistical significance levels are shown in the color bar and gene counts are indicated by the marker size. c. Box plots of the signature score using all genes in the CAIRO HB high-risk rC2 (CLASSES_UP) or low-risk rC1 (CLASSES_DN) gene sets for all nine HB tumors. d. Box plots of the high-risk rC2 or low-risk rC1 gene sets for the five HB tumor cell types.

**Supplemental Figure 9. Sub-clustering of tumor cells within each HB patient.** a. Sub-clustering and correlation heatmap of tumor cells in each patient. Tumor cell sub-clusters are shown in different colors and correlation coefficients between each tumor cell pair are shown in the color bar. b. Heatmap of the five tumor subtype signature scores in each sub-clusters from the nine HB patients. Scaled signature scores are shown in the color bar.

**Supplemental Figure 10. Analysis of basophils and pro-myelocytes from HB tumors and normal liver.** a. UMAP of all basophils from all nine patients annotated by sample type (left) and bar graph showing the proportional contribution of basophils from each patient to each sample type (right). b. UMAP of all basophils from all nine patients annotated by histological subtype (left) and bar graph showing the proportional contribution of basophils from each histological subtype to each sample type (right). c. Heatmap showing differentially expressed genes between basophils found in the adjacent normal tissue versus tumor tissue. d. UMAP of all pro-myelocytes from all nine patients annotated by sample type (left) and bar graph showing the proportional contribution of pro-myelocytes from each patient to each sample type (right). e. UMAP of all pro-myelocytes from all nine patients annotated by histological subtype (left) and bar graph showing the proportional contribution of pro-myelocytes from each histological subtype to each sample type (right). f. Heatmap showing differentially expressed genes between pro-myelocytes found in the adjacent normal tissue versus tumor tissue.

**Supplemental Figure 11. Pseudotime analysis of HB associated erythroid.** a. Left panel, UMAP of HB-associated erythroid cells annotated by the three developmental stages. Right panel, stacked bar charts of HB-associated erythroid in patients 2, 5 and 6 compared with the fetal lever samples from the Popescu. et al. study. b. Pseudotime trajectory of fetal liver erythroid and clustering by the three developmental stages. c. Pseudotime trajectory of HB-associated erythroid cells by the three developmental stages and by patients.

**Supplemental Figure 12. Integrated analysis of tumor cells and fetal liver hepatoblasts.** a. Integrated UMAP of tumor cells with fetal liver hepatoblasts. B. Heatmap of top 10 most differentially expressed genes in each tumor cell cluster and reference fetal liver hepatoblasts. Scaled expressions are shown in the color bar.

**Supplemental Figure 13. Erythroid marker expressions in tumor cells of three patients.** a. Violin plots of seven canonical erythroid markers in all cell types in Patients 2,5 and 6 after ambient RNA decontamination. b. Supervised heatmap showing the expression of canonical erythroid markers and representative markers of three erythroid developmental stages in Patients 2, 5 and 6 tumor cells. c. Representative ligand-receptor interactions between TAMs and tumor cells for patients 2, 5, 6 and other patients. Mean expressions of ligand and receptor pairs are shown in the color bar. Statistical significance levels are indicated by the marker size.

**Supplemental Figure 14. Patient-derived-spheroid morphology and proliferation rate over passages. a.** Brightfield images of patients 2, 6, 7 and 8 tumor spheroids at late passages. **b.** PDS doubling time calculated over five different passages. **c.** Brightfield images of nearby population of patients 7 and 9 cultivated cells, identified as fibroblast-like cells and BEC-like cells. Scale bar = 100 μm.

**Supplemental Figure 15. Broad comparison of freshly isolated tumor cells and patient-derived-spheroids, and identification of spheroid parent cells.** a. UMAP projection of merged scRNA-seq data of cells from tumor tissue and PDS at early and late passage for patients 2, 6, 7, 8 and 9. b.-f. (left panel) UMAPs projection colored by origin (spheroids or tumor) or by cluster annotation (generated from initial annotation) (Figure 1). b.-f. (right panel) Heatmap illustrating signature scores of every cluster generated from signatures of cells from tumor tissue clusters (b. patient 2, c. patient 6, d. patient 7, e. patient 8, f. patient 9).

**Supplemental Figure 16. Identification of a specific subpopulation of patient-derived-spheroid parent cells.** a. c. UMAP projection of freshly isolated tumor cells and spheroids at early passage colored by sub clusters. b. d. Signature score of PDS subclusters calculated with freshly isolated tumor cell cluster signatures. (a., b. patient 6 and c., d. patient 8).

**Supplemental Figure 17. Expression of genes involved in drug metabolism and efflux in PDS.** Dot plots of expression of genes involved in drug metabolism and efflux and corresponding module score calculated for each patient. a. Platinum-based compounds resistance, b. Etoposide resistance, c. Vincristine resistance, d. Verteporfin resistance, e. 5-Fluorouracil resistance, and f. DNA-repair.

## TABLES

**Supplemental Table 1. De-identified clinical information of the nine HB patients in the study.**

**Supplemental Table 2. Single-cell RNA sequencing cell quality summary and annotation information.** Numbers of cells captured, mean and median numbers of genes/UMIs, and mean percentage of mitochondrial gene expressions are shown for each sequenced array. Manual annotations are validated and consistent with automated annotation by SingleR. Signature gene sets generated from manual annotation are specifically listed.

**Supplemental Table 3. Top 50 differemtially expressed genes in each tumor cell cluster.** Gene symbols, average expression log2 fold change, and adjusted p-values are shown.

**Supplemental Table 4. Signature gene sets of the five HB tumor subtypes. Gene symbols, average expression log2 fold change, and adjusted p-values are shown.**

**Supplemental Table 5. Origin of cultivated Patient-derived spheroids (Freshly isolated cells or Cryopreserved freshly isolated cells) and ability of cells to grow at short term (2-4 passages) or long term (more than 16 passages).**

