## Supplementary figures and images for "Single-cell analysis of hepatoblastoma identifies distinct tumor cell signatures that predict susceptibility to chemotherapy using patient-specific tumor spheroids"

### Supplemental Figure 1

Supp Figure 1

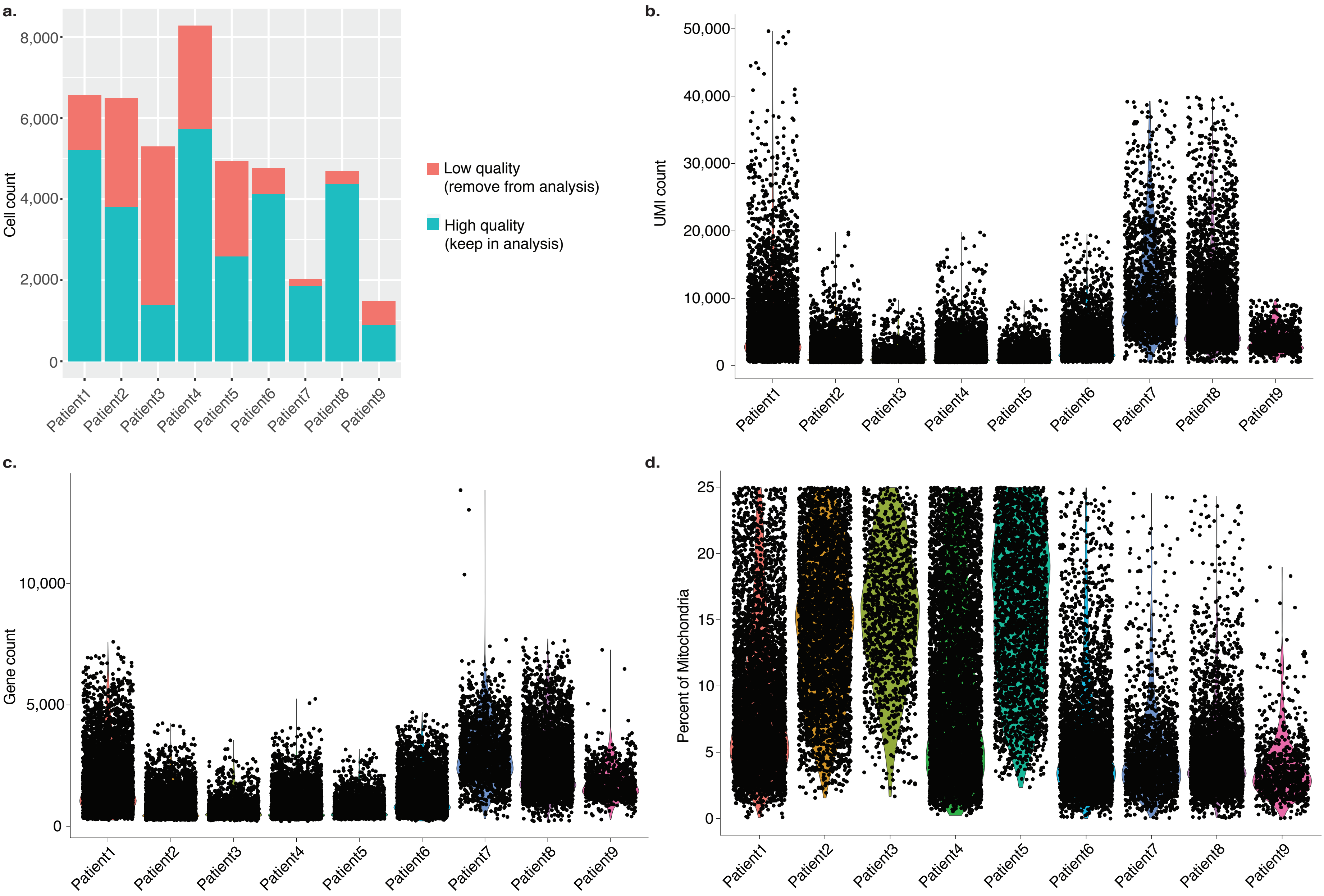

### Supplemental Figure 2

Supp Figure 2

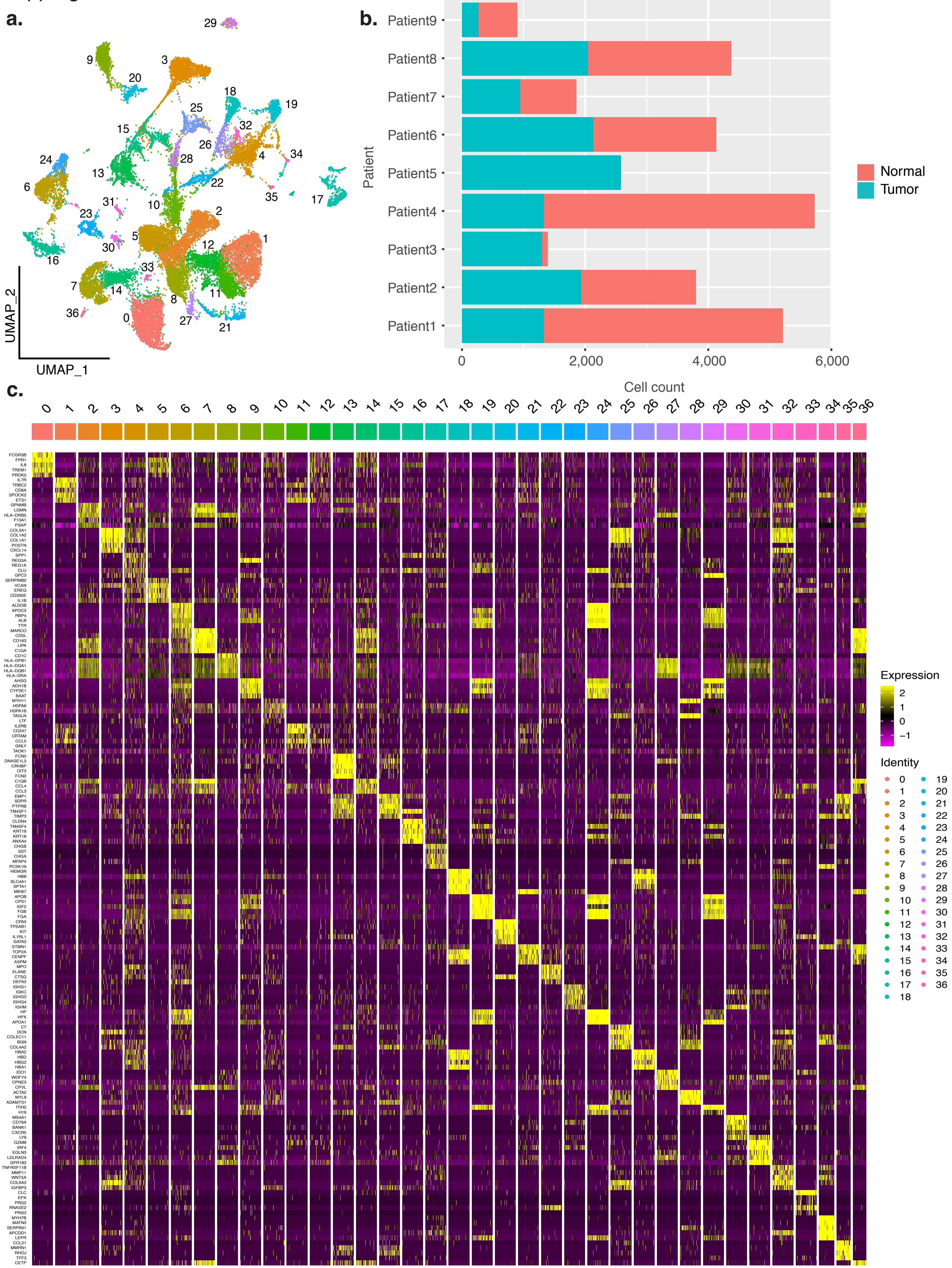

### Supplemental Figure 3

Supp Figure 3

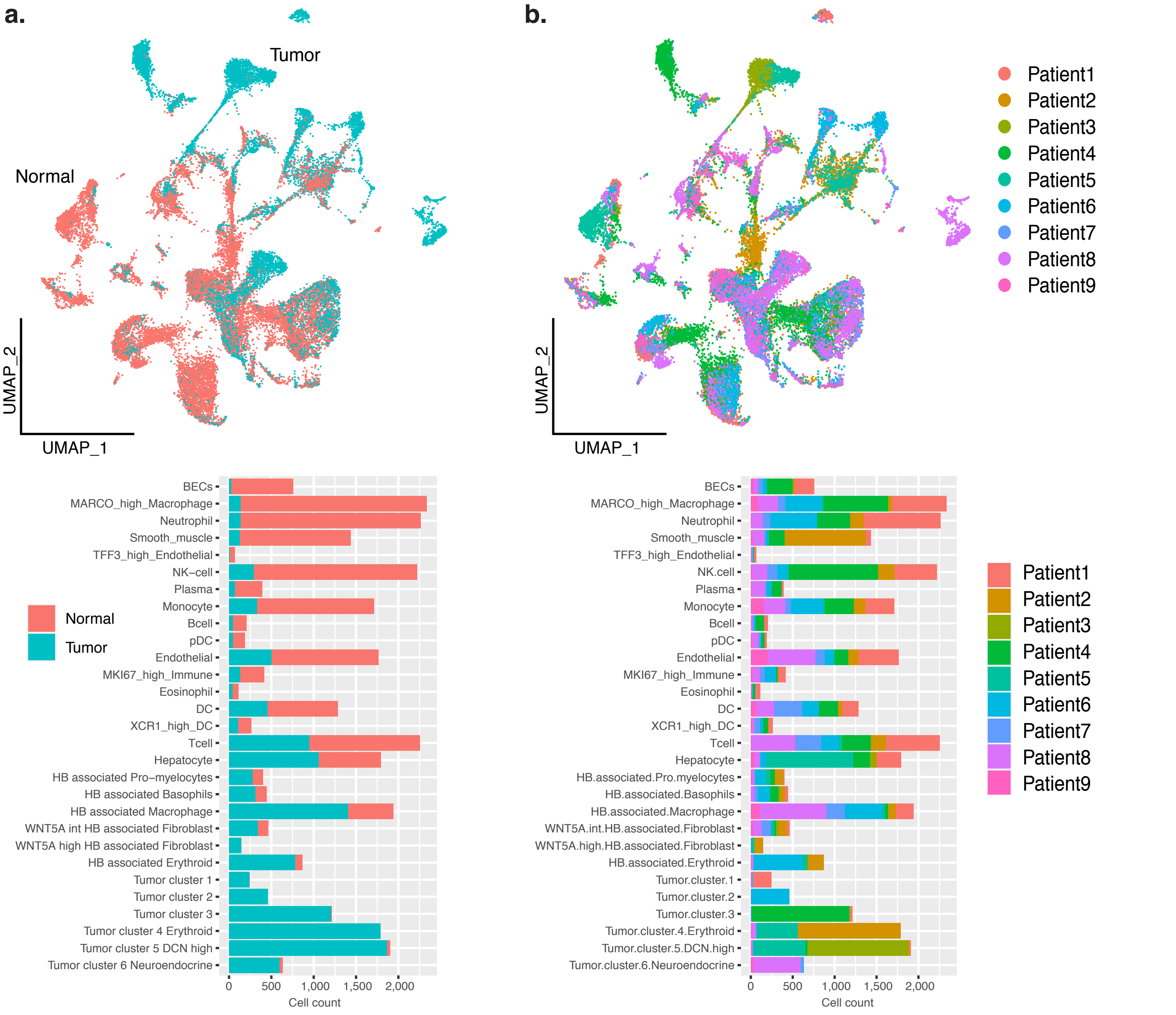

### Supplemental Figure 4

Supp Figure 4

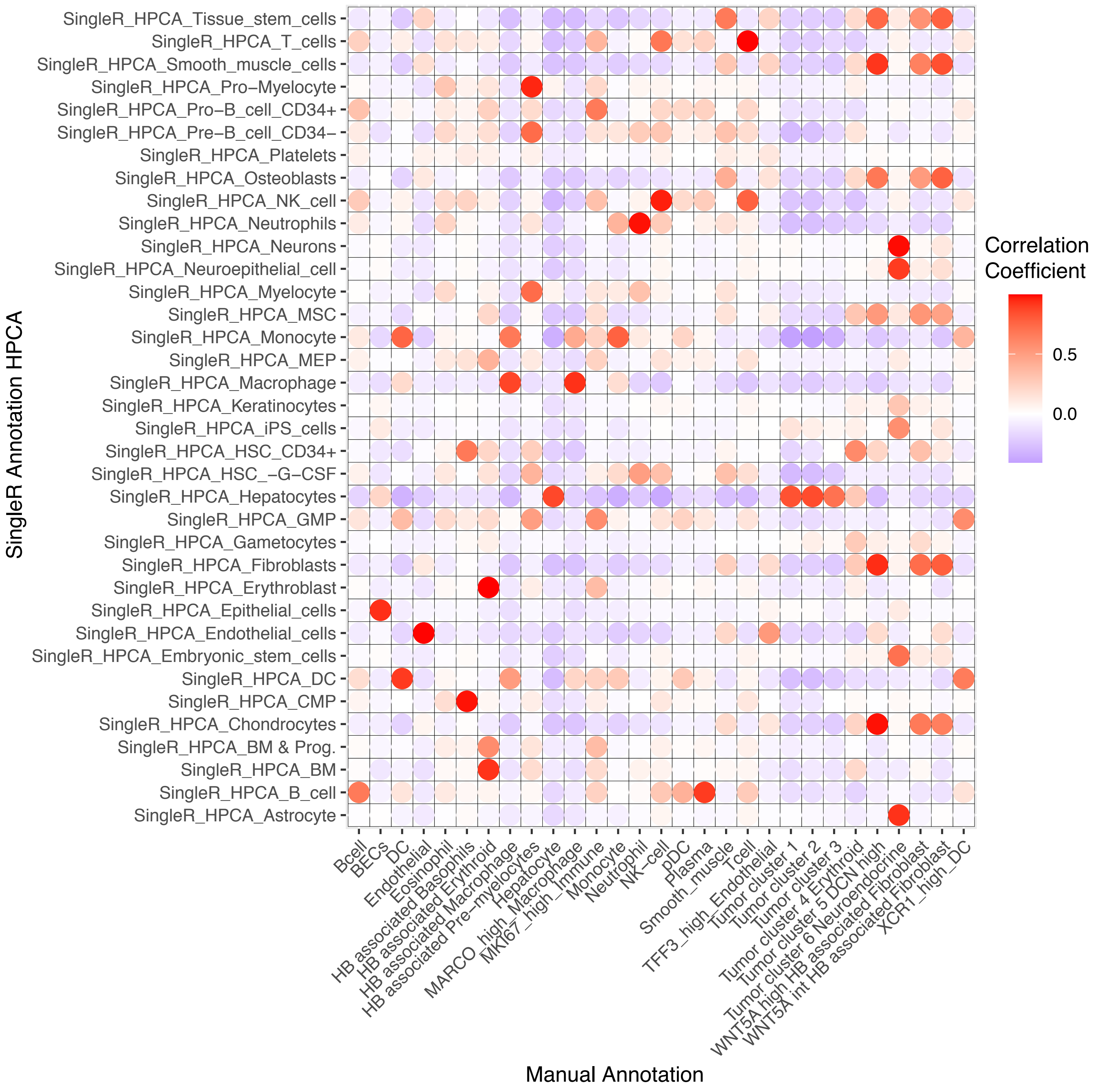

### Supplemental Figure 5

Supp Figure 5

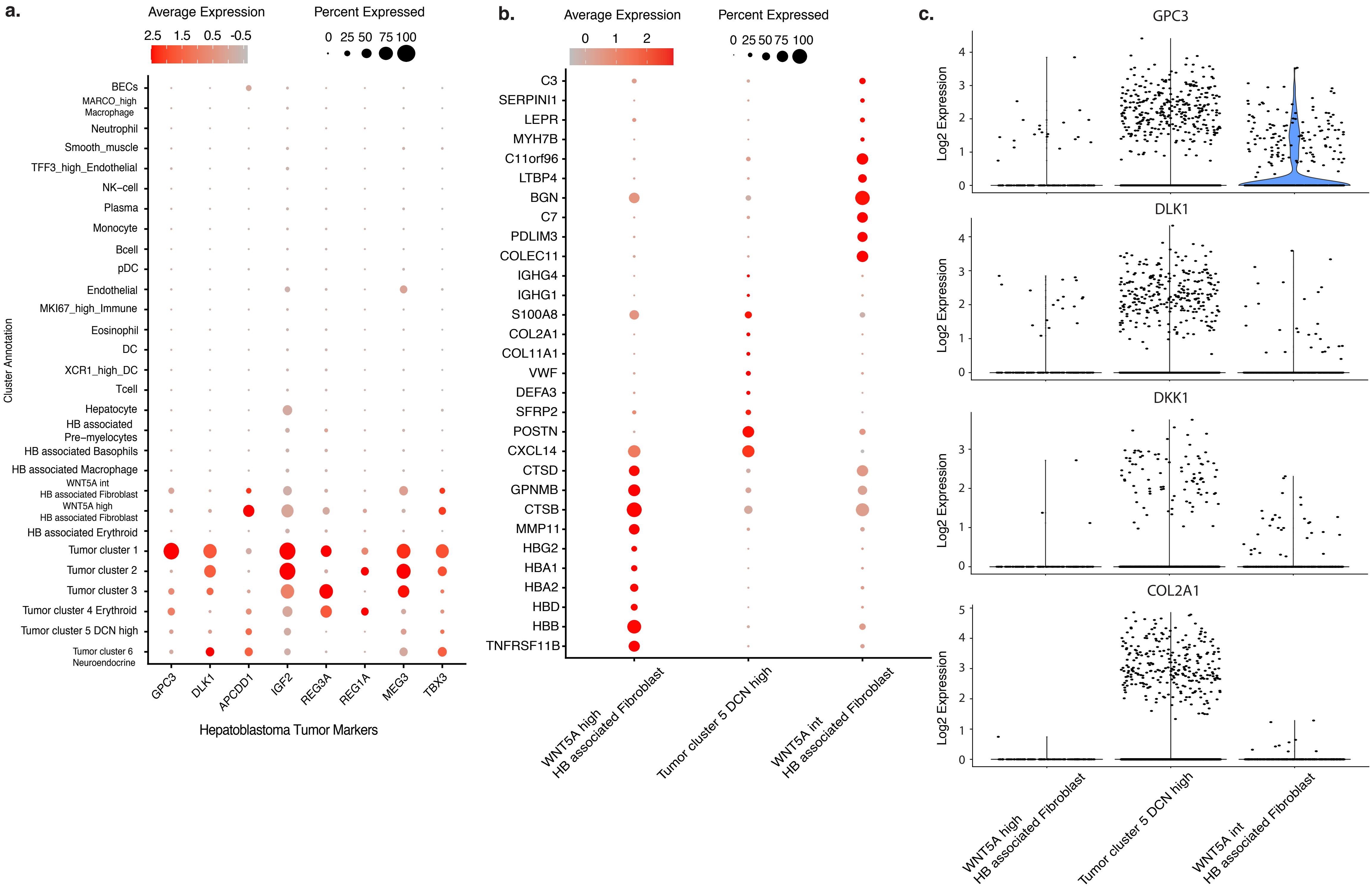

### Supplemental Figure 6

Supp Figure 6

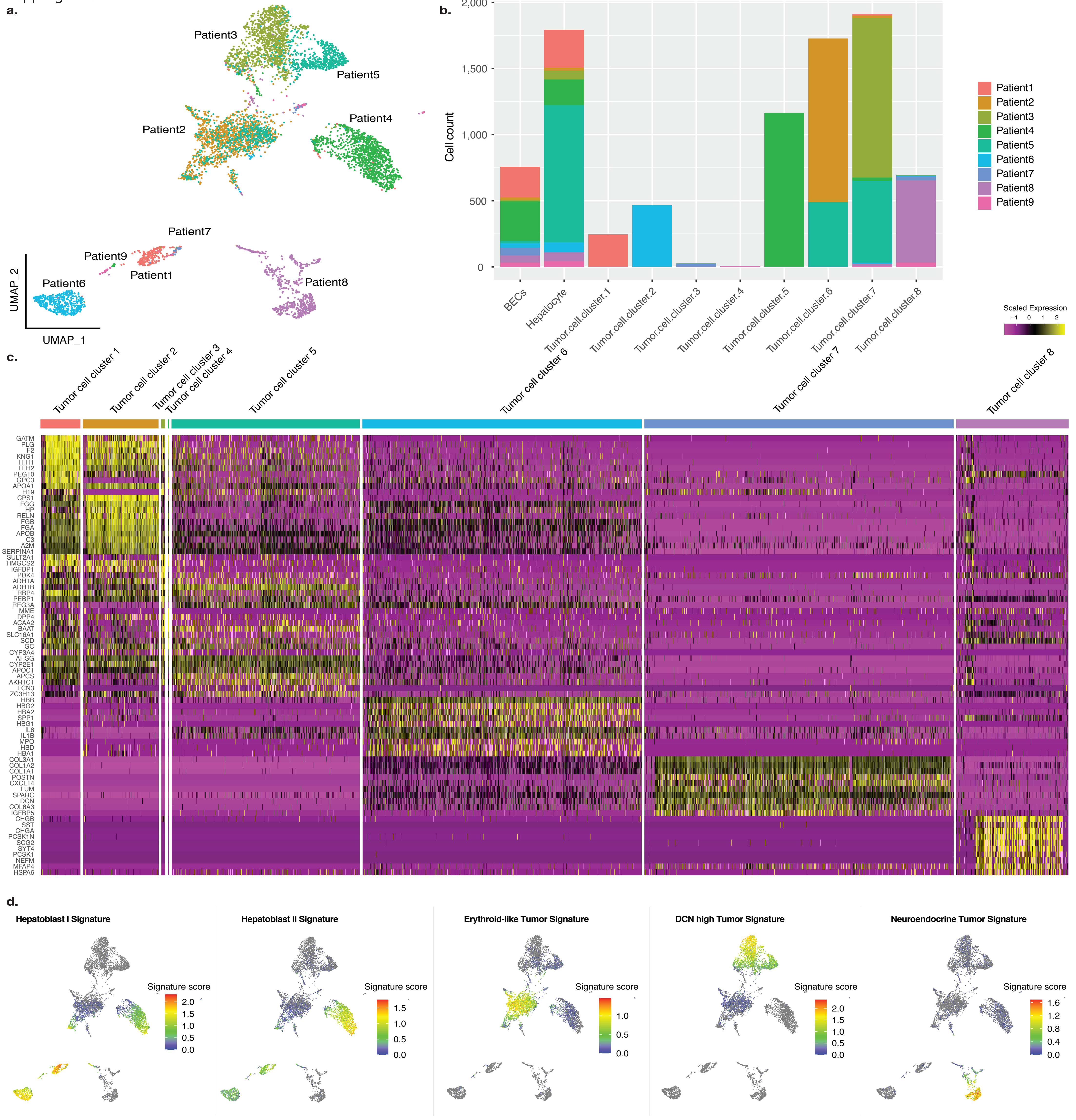

### Supplemental Figure 7

Supp Figure 7

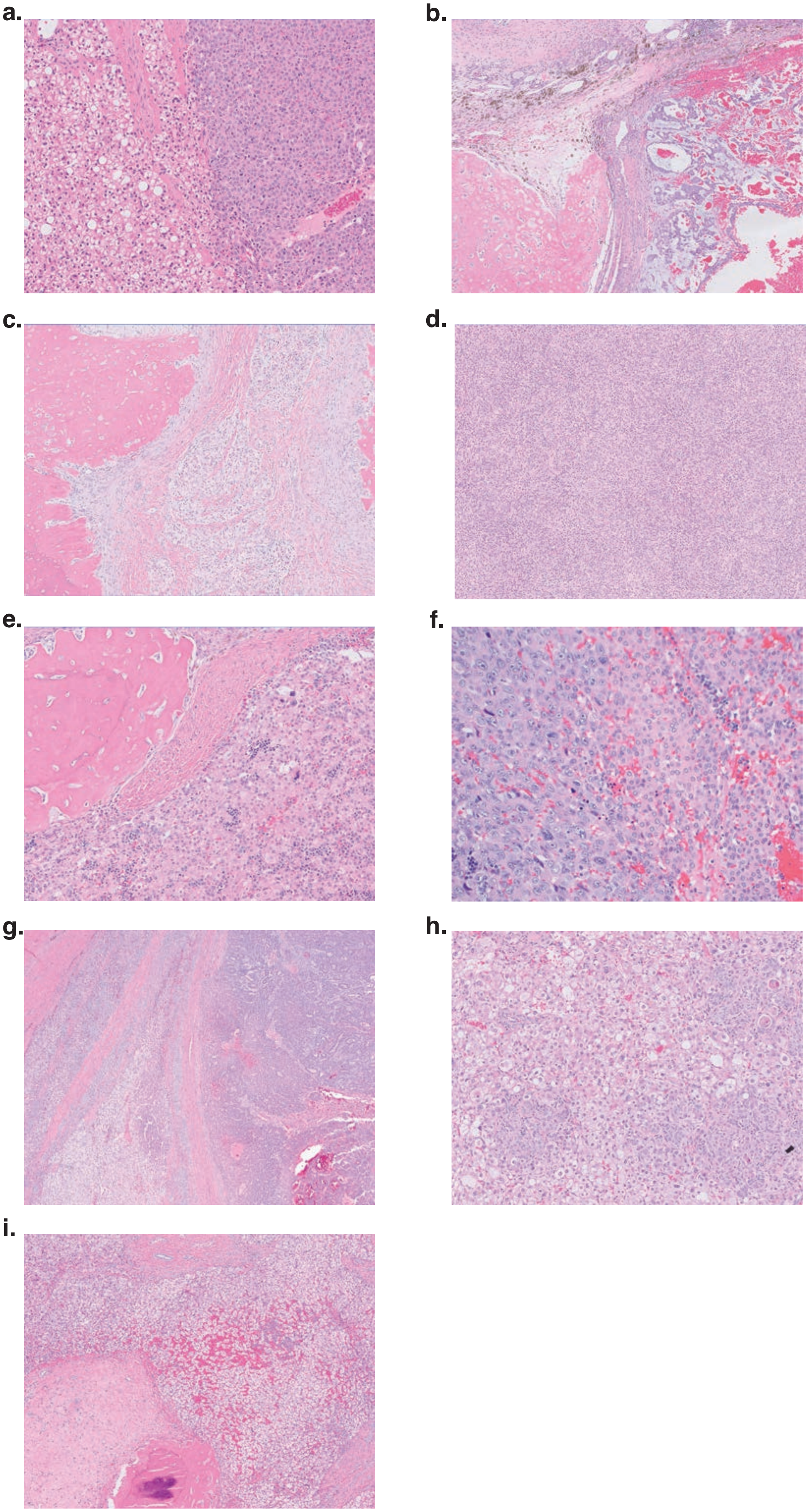

### Supplemental Figure 8

Supp Figure 8

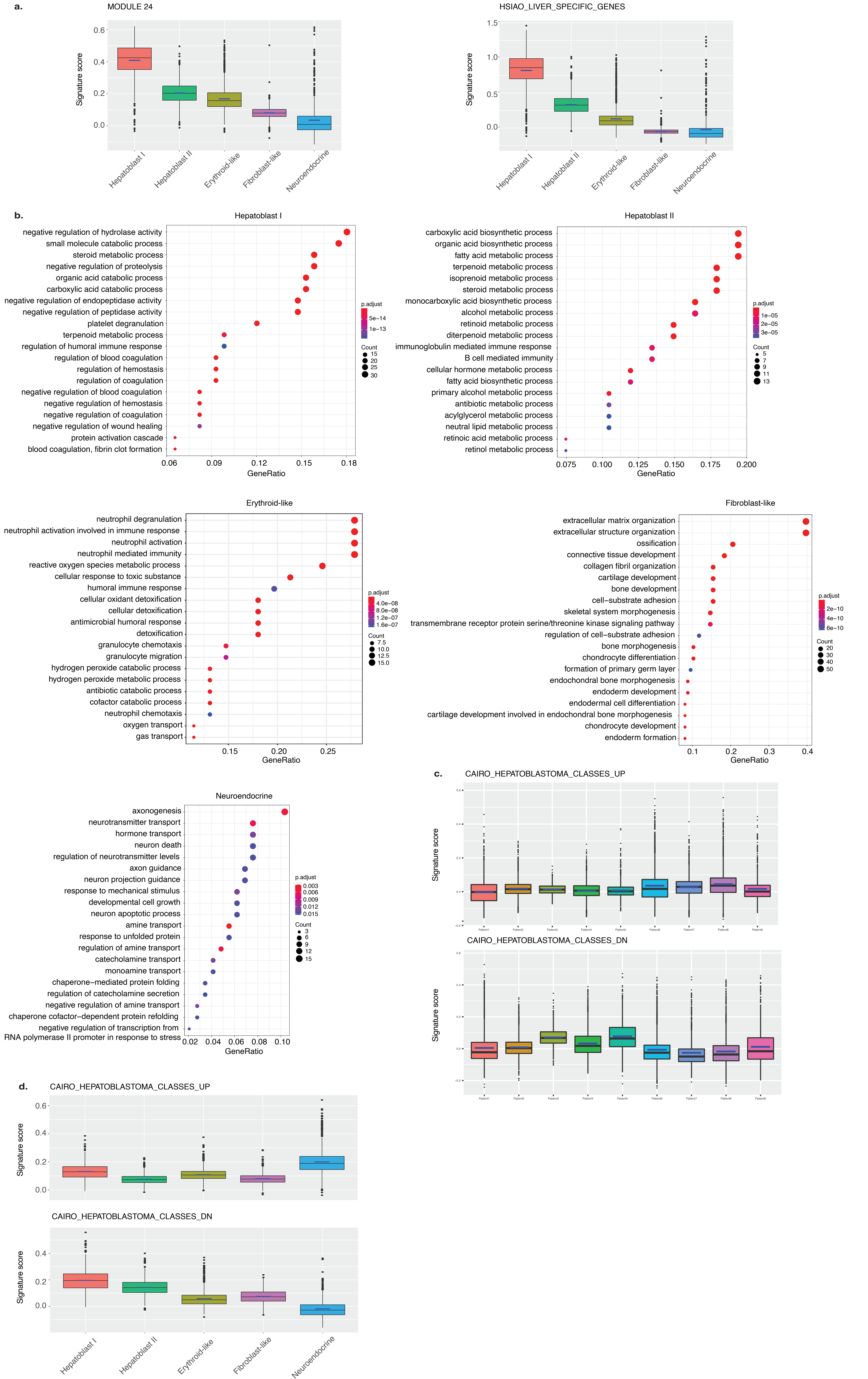

### Supplemental Figure 9

Supp Figure 9

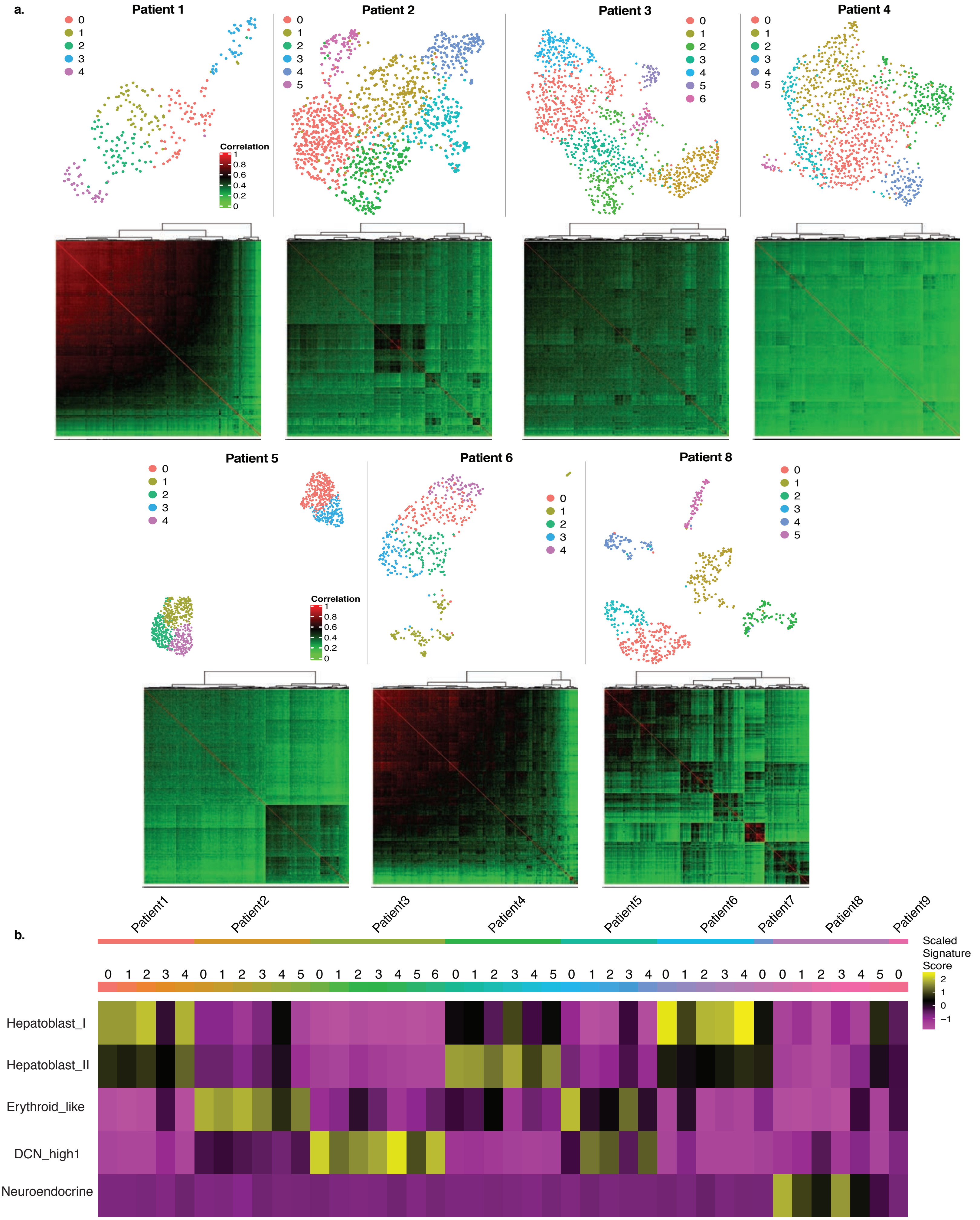

### Supplemental Figure 10

Supp Figure 10

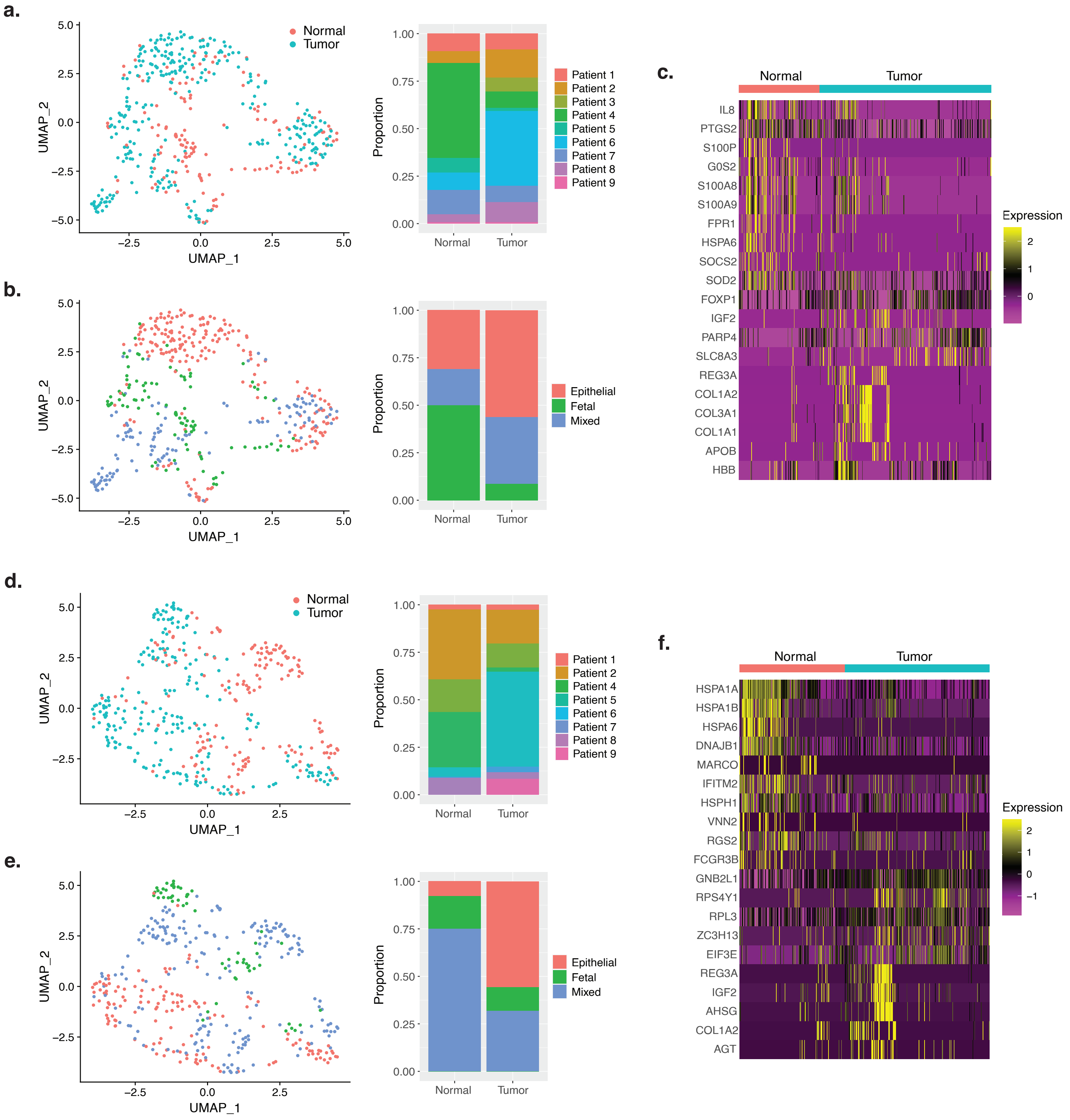

### Supplemental Figure 11

Supp Figure 11

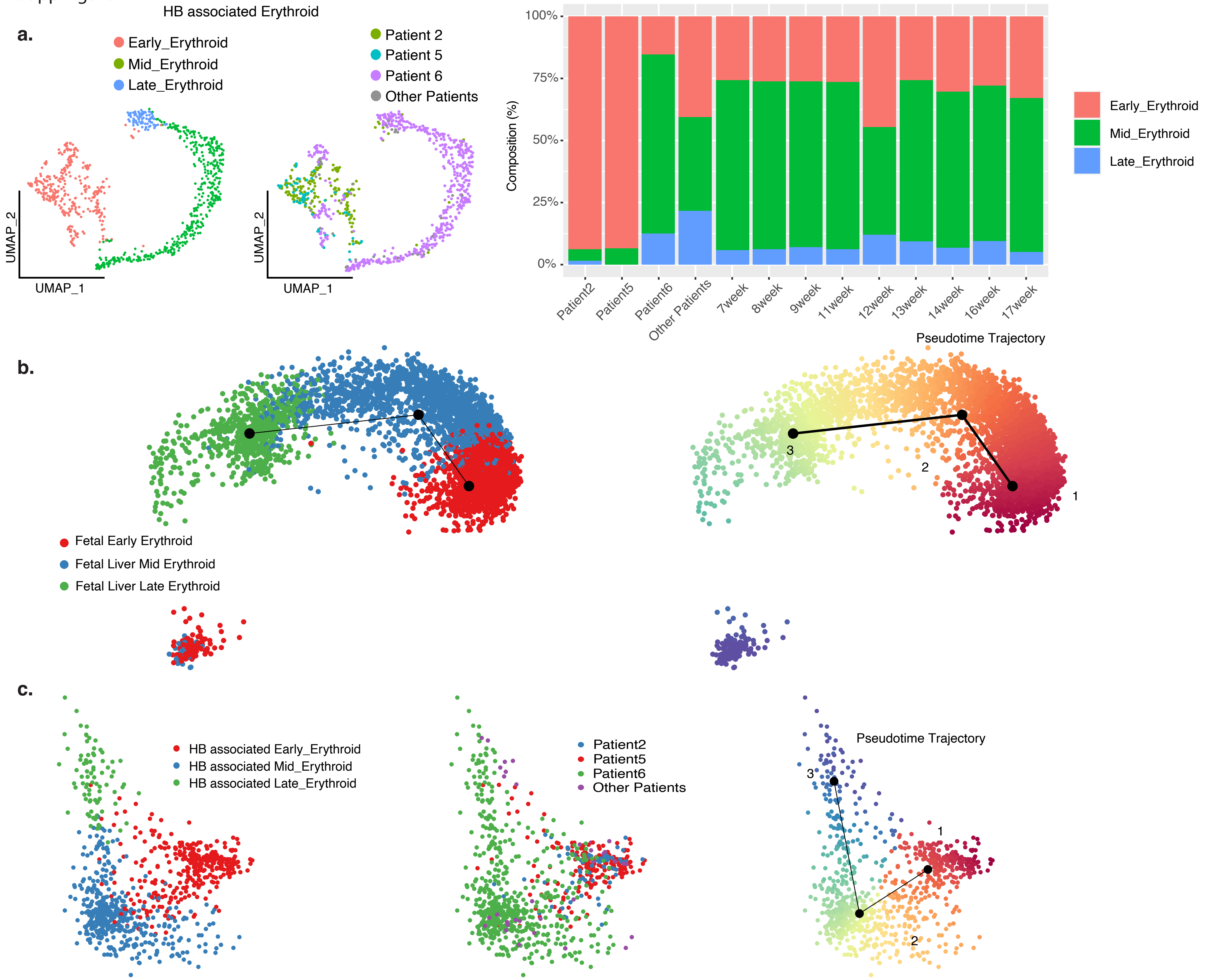

### Supplemental Figure 12

Supp Figure 12

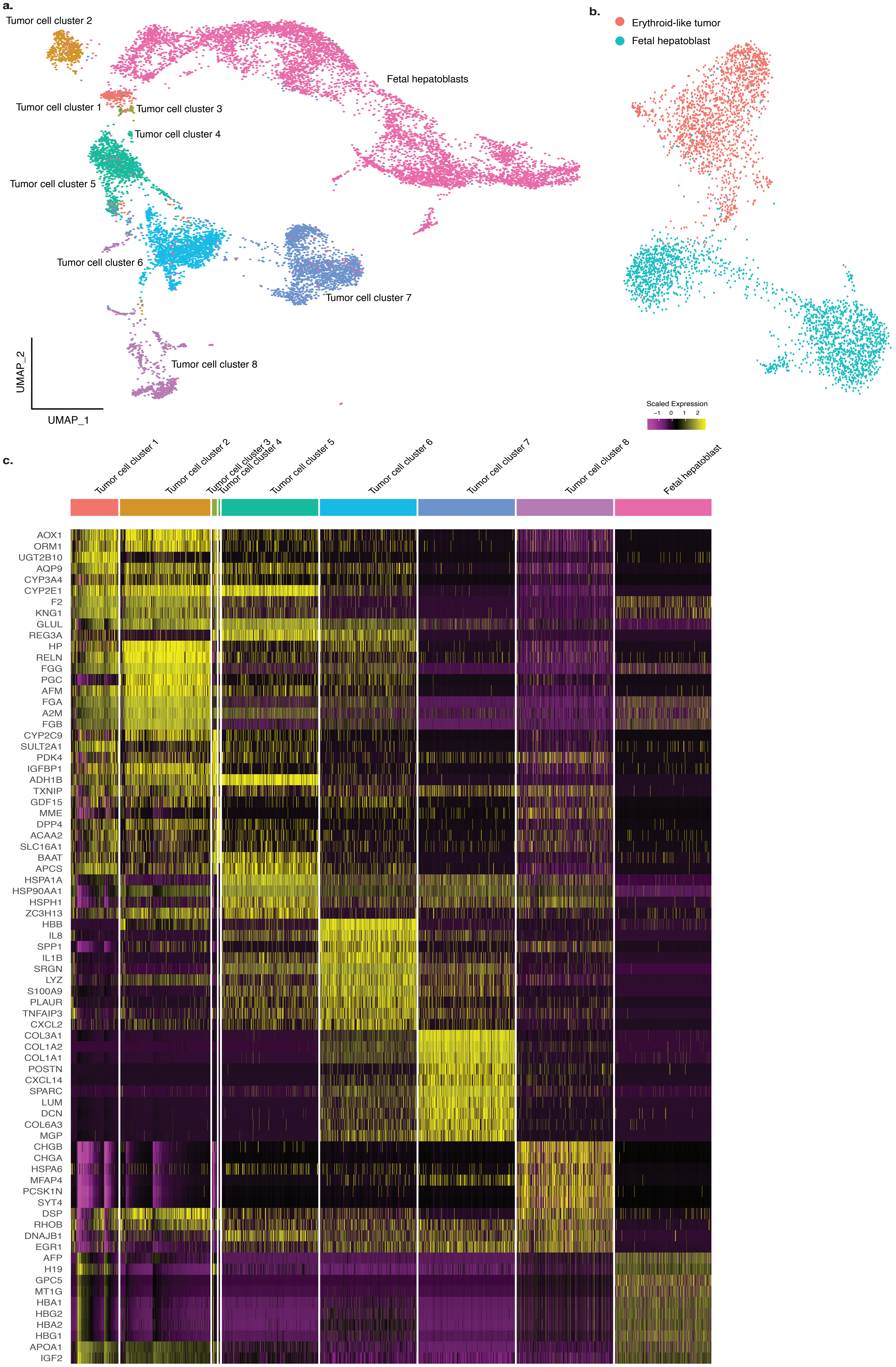

### Supplemental Figure 13

Supp Figure 13

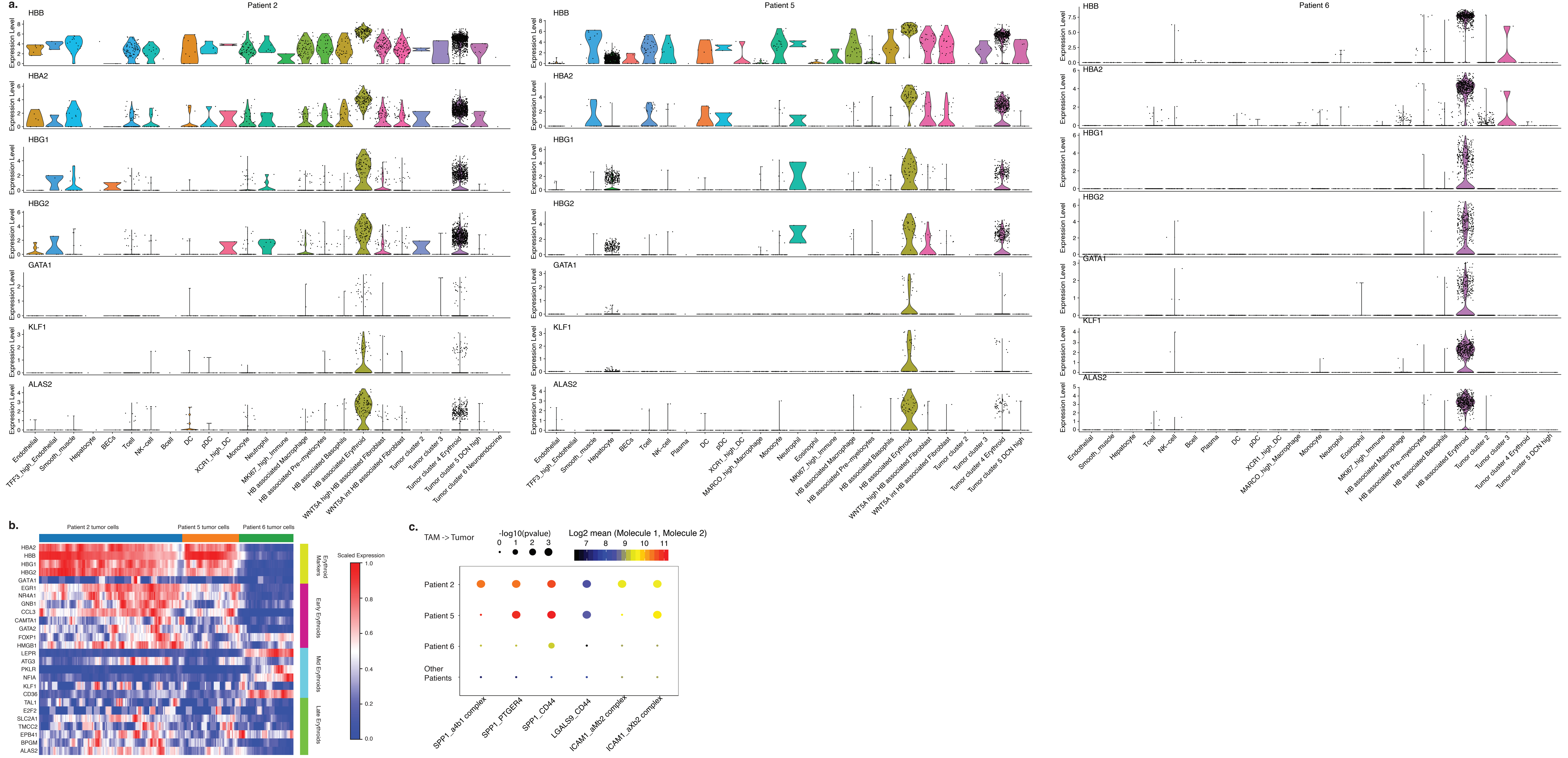

### Supplemental Figure 14

Supp Figure 14

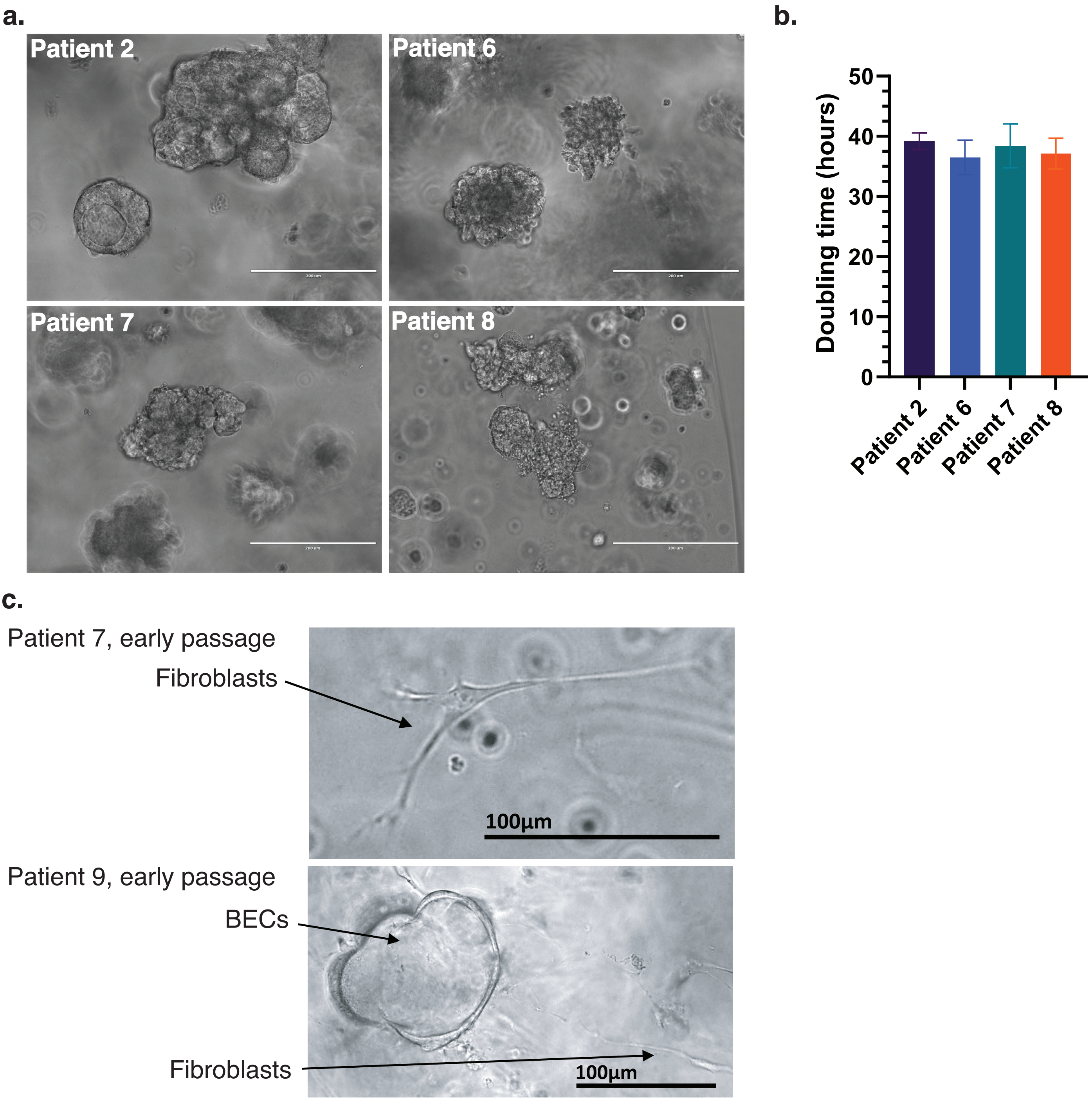

### Supplemental Figure 15

Supp Figure 15

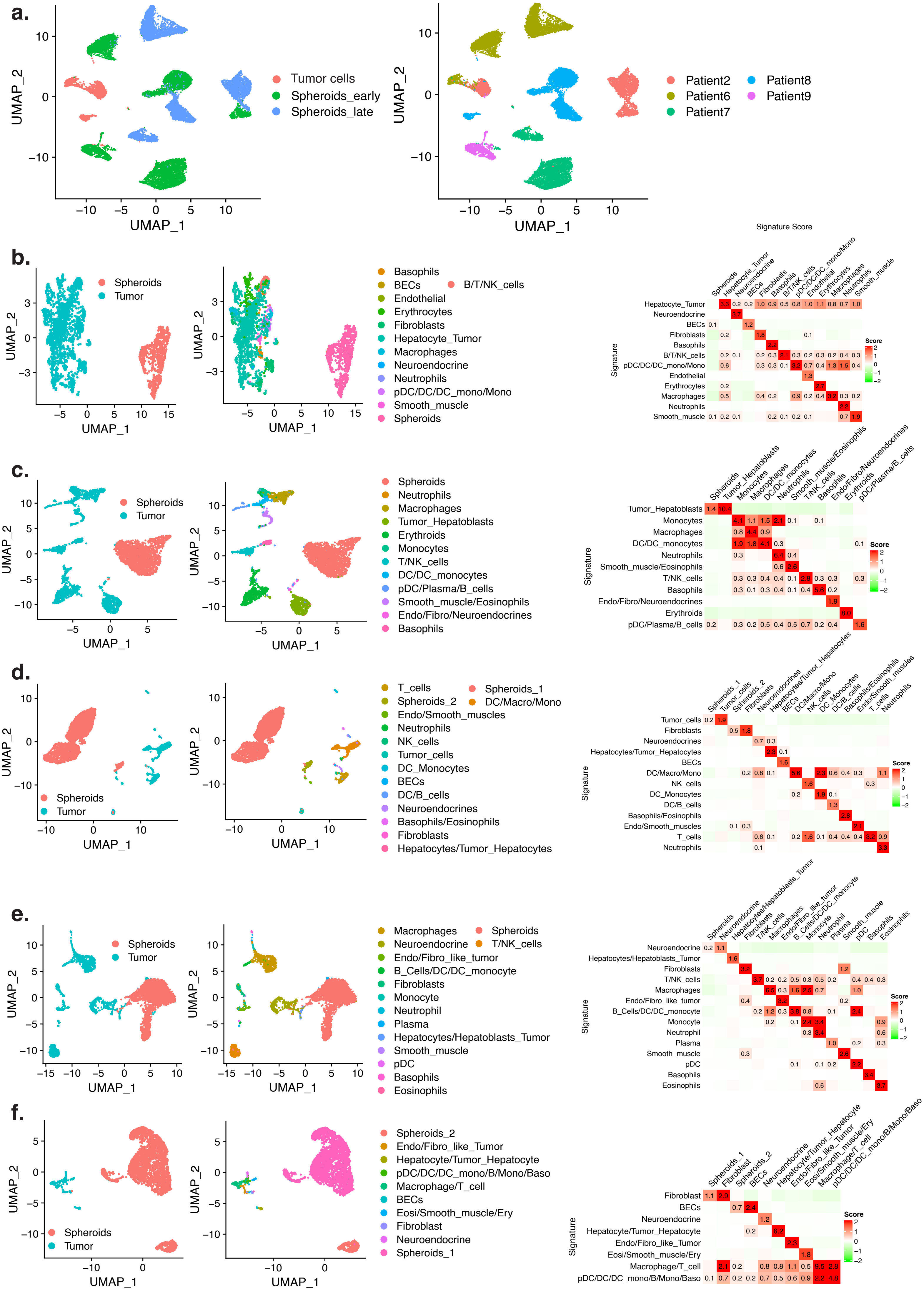

### Supplemental Figure 16

Supp Figure 16

**a.**

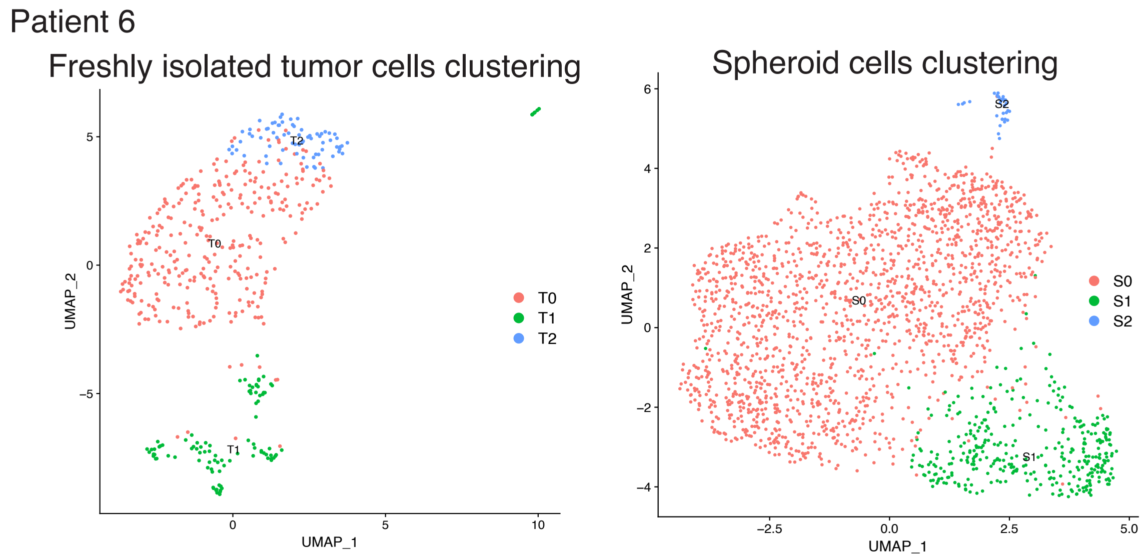


**b.**

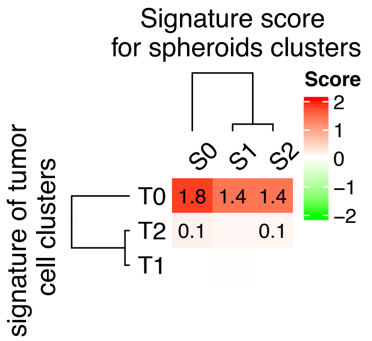


**c.**

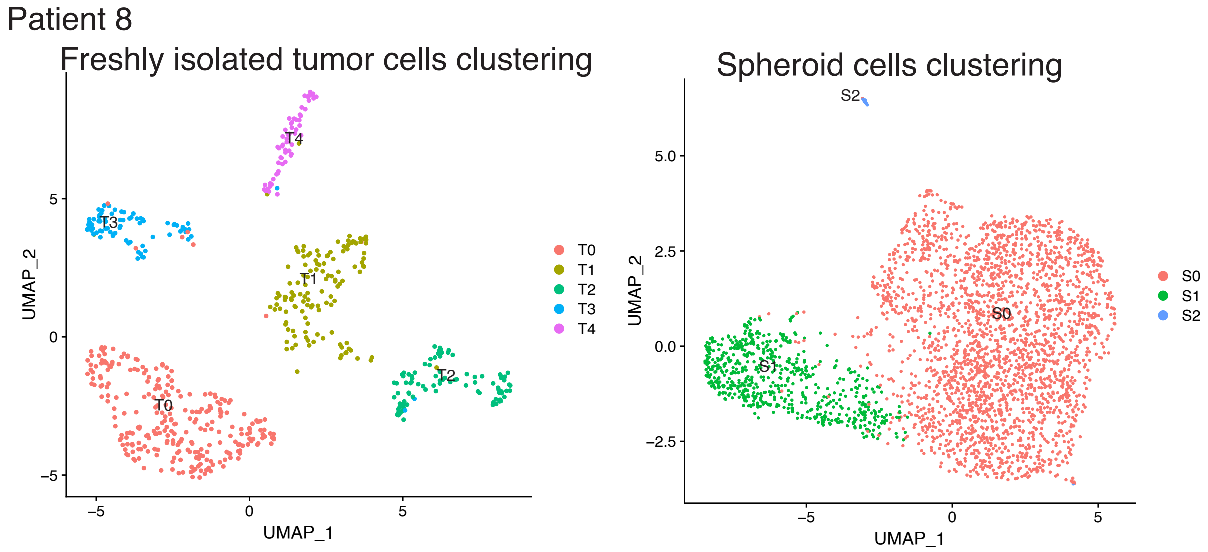


**d.**

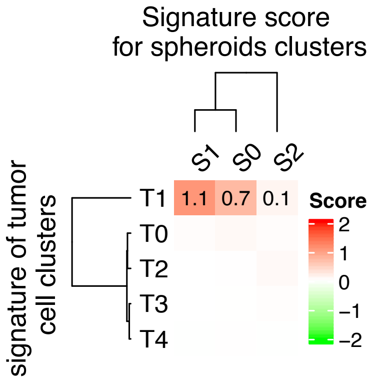

### Supplemental Figure 17

Supp Figure 17

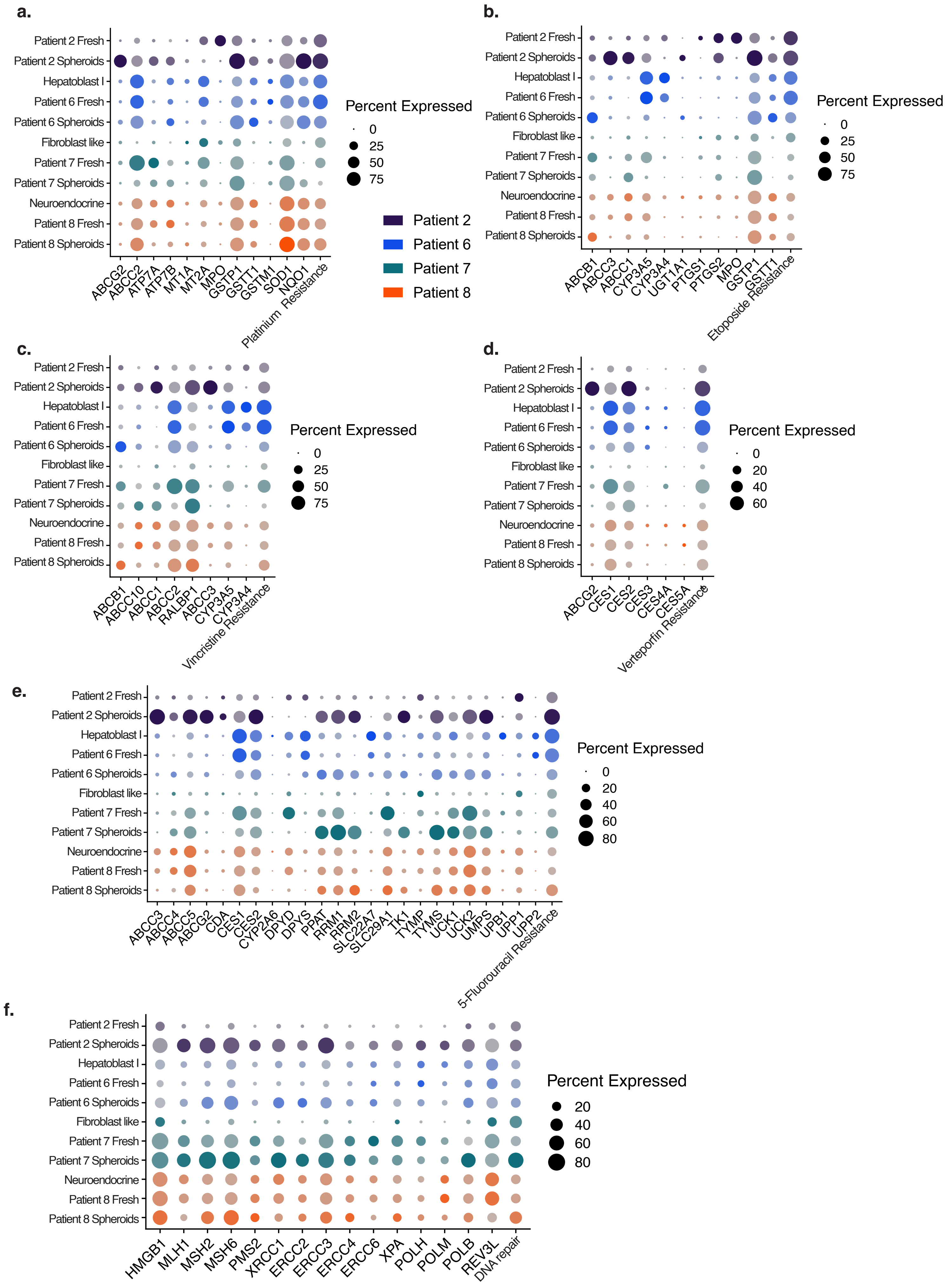
